# Mobile barrier mechanisms for Na^+^-coupled symport in an MFS sugar transporter

**DOI:** 10.1101/2023.09.18.558283

**Authors:** Parameswaran Hariharan, Yuqi Shi, Satoshi Katsube, Katleen Willibal, Nathan D. Burrows, Patrick Mitchell, Amirhossein Bakhtiiari, Samantha Stanfield, Els Pardon, H. Ronald Kaback, Ruibin Liang, Jan Steyaert, Rosa Viner, Lan Guan

## Abstract

While many 3D structures of cation-coupled transporters have been determined, the mechanistic details governing the obligatory coupling and functional regulations still remain elusive. The bacterial melibiose transporter (MelB) is a prototype of the Na^+^-coupled major facilitator superfamily transporters. With a conformational nanobody (Nb), we determined a low-sugar affinity inward-facing Na^+^-bound cryoEM structure. Collectively with the available outward-facing sugar-bound structures, both the outer and inner barriers were localized. The N-and C-terminal residues of the inner barrier contribute to the sugar selectivity pocket. When the inner barrier is broken as shown in the inward-open conformation, the sugar selectivity pocket is also broken. The binding assays by isothermal titration calorimetry revealed that this inward-facing conformation trapped by the conformation-selective Nb exhibited a greatly decreased sugar-binding affinity, suggesting the mechanisms for the substrate intracellular release and accumulation. While the inner/outer barrier shift directly regulates the sugar-binding affinity, it has little or no effect on the cation binding, which is also supported by molecular dynamics simulations. Furthermore, the use of this Nb in combination with the hydron/deuterium exchange mass spectrometry allowed us to identify dynamic regions; some regions are involved in the functionally important inner barrier-specific salt-bridge network, which indicates their critical roles in the barrier switching mechanisms for transport. These complementary results provided structural and dynamic insights into the mobile barrier mechanism for cation-coupled symport.

## Introduction

Cation-coupled transporters play critical roles in many aspects of cell physiology and pharmacokinetics and are involved in many serious diseases including cancers, metabolic diseases, neurodegenerative diseases, etc. Their specific locations and functions make them viable drug targets^1,2^. Following decades of effort, the structures of many transporters have been determined by x-ray crystallography and, more recently, by cryoEM^3–14^. However, we still do not have a deep understanding of the structural basis for substrate accumulation to high concentrations, the mechanisms for the obligatorily coupled transport, as well as the cellular regulation of transporters’ activities to meet metabolic needs.

*Salmonella enterica* serovar Typhimurium melibiose permease (MelB_St_) is the prototype Na^+^/solute transporter of the major facilitator superfamilies (MFS)^11,15–24^. It catalyzes the stoichiometric symport of galactopyranoside with Na^+^, H^+^, or Li^+^^25^. Crystal structures of sugar-bound states have been determined, which reveals the sugar recognition mechanism^19^. In all solved x-ray crystal structures, MelB_St_ is in an outward-facing conformation; the observed physical barrier of the sugar translocation, named the inner barrier, prevents the substrate from moving into the cytoplasm directly. It is well-known that transporters change their conformation to allow the substrate translocation from one side of the membrane to the other and conformational dynamics is the intrinsic feature necessary for their functions^3,6,7,12,13,26,27^.

To understand how the sugar substrate and the coupling cations dictate transporter function, we have established thermodynamic binding cycles in MelB_St_ using isothermal titration calorimetry (ITC) measurements and identified that the positive cooperativity of the binding of cation and melibiose as the core symport mechanism^17^; furthermore, the binding of the second substrate changes the thermodynamic feature of heat capacity change from positive to negative, suggesting primary dehydration processes when both substrates concurrently occupy the transporter^28^. While the structural origins have not yet been completely determined yet, they may be related to forming an occluded intermediate conformation, a necessary state for a global conformational transition.

To determine structurally unresolved states of MelB_St_, nanobodies (Nbs) were raised against the wild-type MelB_St_. One group of Nbs (Nb714, Nb725, and Nb733) has been identified to interact with the cytoplasmic side of MelB_St_ and their binding completely inhibits MelB_St_ transport activity^29^. MelB_St_ bound with Nb725 or Nb733 retained its binding to Na^+^, but its affinity for melibiose is inhibited. The MelB_St_/Nb733 complex also retained the affinity to its physiological regulator dephosphorylated EIIA^Glc^ of the glucose-specific phosphoenolpyruvate:phosphotransferase system (PTS). EIIA^Glc^, the central metabolite regulator in bacterial catabolic repression^30–32^, greatly inhibited the galactoside binding to non-PTS permeases MelB_St_^33^ and the H^+^-coupled lactose permease (LacY)^34^, but still retained the binding of Na^+^ to MelB_St_^29^. Thus, the functional modulation of the two Nbs on MelB_St_ mimics the binding of EIIA^Glc^. The crystal structure of dephosphorylated EIIA^Glc^ binding to the maltose permease MalFGK_2_, an ATP-binding cassette transporter, has been determined^35^, but there is no structural information on EIIA^Glc^ binding to the MFS transporters. Determination of the structures of MelB_St_ in complex with this group of Nbs is important and will reveal critical information to understand the manner by which EIIA^Glc^ regulates the large group of MFS sugar transporters.

In the current study, we applied cryoEM single-particle analysis (cryoEM-SPA) and determined the near-atomic resolution structure of MelB_St_ complexed with a Nb725_4, a modified Nb725 with a designed scaffold which enables NabFab binding^36^. The use of the Nb provided a Na^+^-bound lower sugar-affinity inward-facing conformation of MelB_St_. Furthermore, our studies using this conformation-selective Nb in combination with hydrogen/deuterium exchange mass spectrometry (HDX-MS) demonstrate a powerful approach for characterizing protein conformational flexibility^37,38^, and also provides critical dynamic information to understand how the substrate binding drives the conformational transition necessary for the substrate translocation in MelB_St_.

## Results

### CDR grafting to generate a hybrid Nb and its functional characterization

To facilitate structure determination of MelB_St_ complexed with the conformation selective Nb by cryoEM-SPA, a novel universal tool based on a NabFab was applied to increase the effective mass of small particles^36^, since MelB_St_ (mass, 53.5 kDa) has limited extramembrane mass. A hybrid Nb725_4 was created by complementarity-determining region (CDR) grafting or antibody reshaping^39^ to transfer the binding specificity of MelB_St_ Nb725 to the TC-Nb4 recognized by the NabFab^36^ (**sFig. 1a**). Extensive *in vivo* and *in vitro* functional characterization revealed that the hybrid Nb725_4 possesses the binding properties of its parents Nbs (Nb725 and TC-Nb41) with slightly compromised affinities to MelB_St_. Nb725_4 binds to MelB_St_ intracellularly and inhibits MelB-mediated melibiose fermentation and active transport, with an equilibrium dissociation constant (*K*_d_) of 3.64 ± 0.62 µM, compared to 1.58 ± 0.42 µM for Nb725 (**Fig. 1a-d**; **Table 1**).

**Fig. 1.**
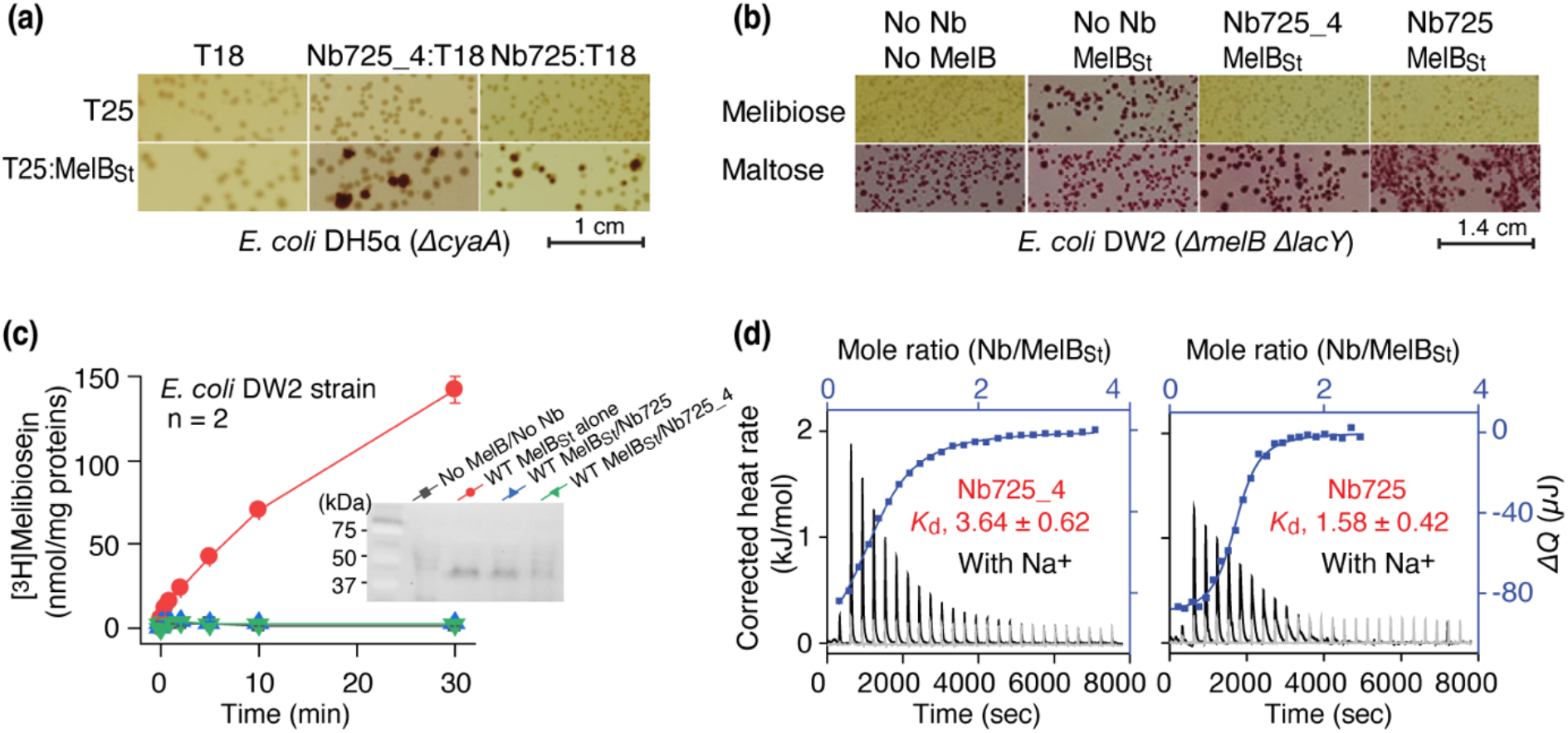
Functional characterizations of the hybrid Nb725_4. **(a)** *In vivo* two-hybrid interaction assay. Two compatible plasmids encoding T25:MelB_St_ and Nb:T18 were transformed into *E. coli* DH5*α cyaA* cells and plated on the maltose-containing MacConkey agar plate as described in Methods. The irregular red colonies are typical of a positive two-hybrid test indicating protein-protein interactions. **(b)** Nb inhibition of melibiose fermentation. Two compatible plasmids encoding MelB_St_ and Nb725 or Nb725_4 were transformed into *E. coli* DW2 cells [∆*melB*∆*lacYZ*] and plated on the MacConkey agar plate containing maltose (as the positive control) and melibiose (for testing transport activity of MelB_St_) as the sole carbon source. **(c)** [^3^H]Melibiose transport assay with *E. coli* DW2 cells. The cells transformed by two compatible plasmids encoding the MelB_St_ and Nb725 or Nb725_4 were prepared for [^3^H]melibiose transport assay at 0.4 mM (specific activity of 10 mCi/mmol) and 20 mM Na^+^ as described in Methods. The cells transformed with the two empty plasmids without MelB or Nb were the negative control. **Inset**, Western blot. MelB_St_ expression under the co-expression system was analyzed by isolating the membrane fractions. An Aliquot of 50 μg was loaded on each well and MelB_St_ protein was probed by anti-His antibody. **(d)** Nb binding to MelB_St_ by ITC measurements. As described in Methods, the thermograms were collected with the Nano-ITC device (TA Instrument) at 25 °C. Exothermic thermograms shown as positive peaks were obtained by titrating Nbs (0.3 mM) into the MelB_St_-free buffer (gray) or MelB_St_ (35 mM)-containing buffer (black) in the Sample Cell and plotted using bottom/left (x/y) axes. The binding isotherm and fitting of the mole ratio (Nb/MelB_St_) vs. the total heat change (f..*Q*) using one-site independent-binding model were presented by top/right (x/y) axes. The dissociation constant *K*_d_ was presented at mean ± SEM (number of tests = 6-7).

**Table 1.**
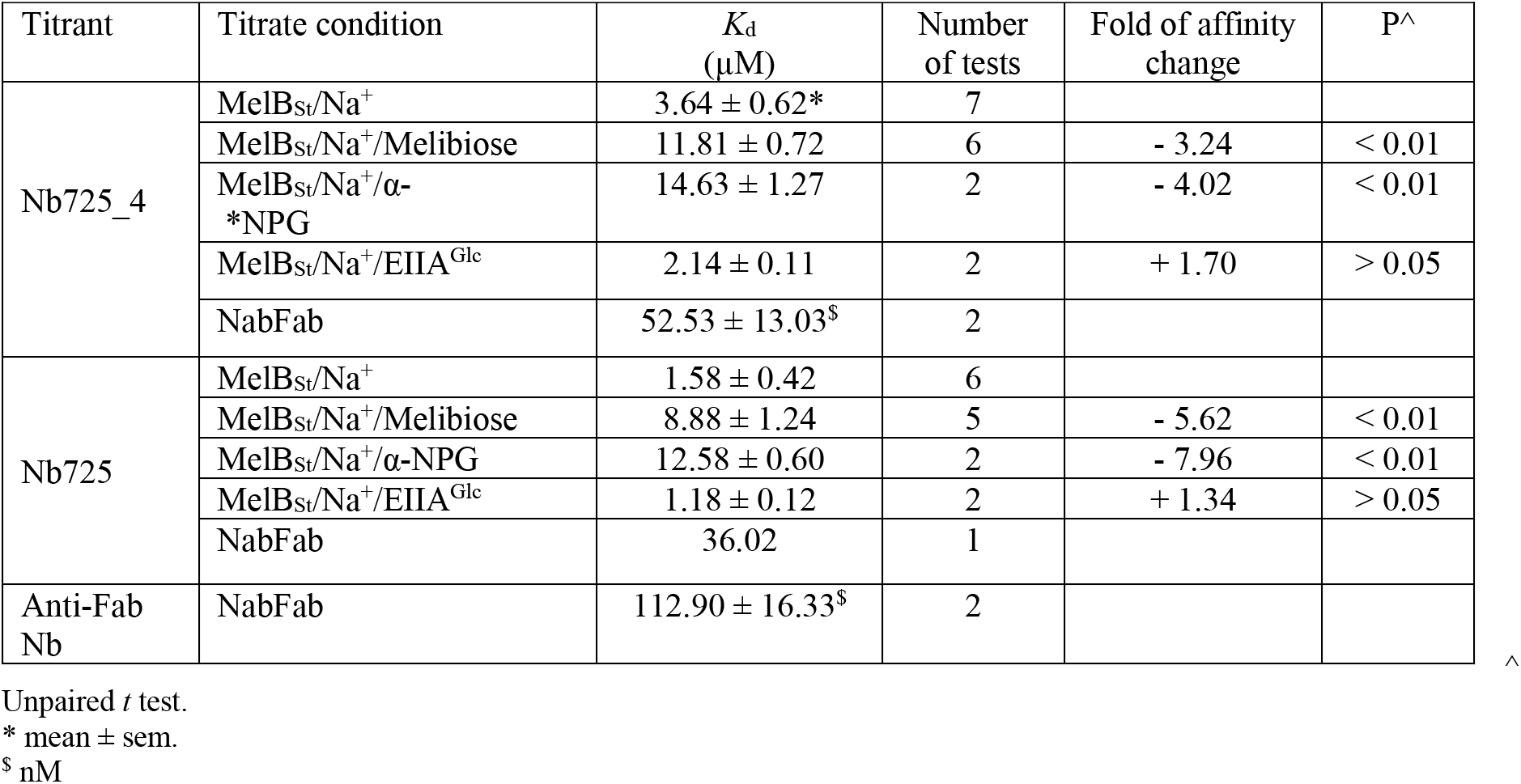
Nbs binding.

The parent Nb725 had a poor binding to the NabFab (**sFig. 1b**)**_,_** but Nb725_4 exhibited greatly increased affinity to NabFab at a *K*_d_ value of 52.53 ± 13.03 nM (**Fig. 1c**; **Table 1**). The anti-Fab Nb binding to NabFab was two-fold poorer than Nb725_4 at a *K*_d_ value of 112.90 ± 16.33 nM (**sFig. 1d**; **Table 1**). The complex containing all four proteins including MelB_St_, Nb725_4, NabFab, and anti-Fab Nb showed shift peak-shift in the gel-filtration chromatography (**sFig. 1e**).

Nb725_4 binding properties were further analyzed in the presence of the MelB ligands. As expected, the presence of melibiose yielded 6-and 3-fold inhibition, and α-NPG afforded 8-and 4-fold inhibition, on the binding of Nb725 and Nb725_4, respectively (**Table 1**; **sFig. 2**). The physiological regulatory protein EIIA^Glc^ (**Table 1**; **sFig. 2**) showed no inhibition on the binding affinity of both Nbs.

**Fig. 2.**
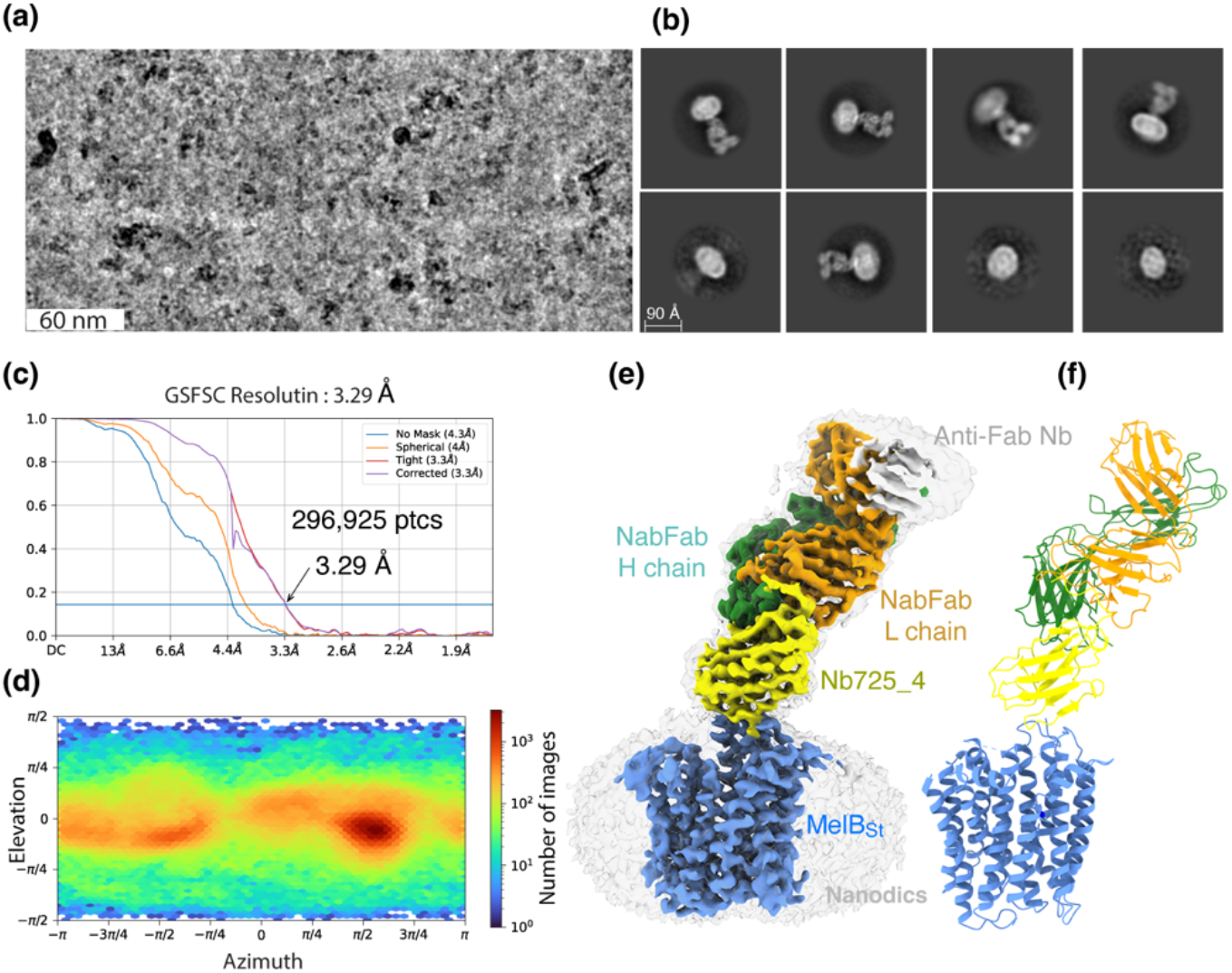
CryoEM-SPA. The sample containing the wild-type MelB_St_ in lipids nanodiscs, the MelB_St_-specific Nb725_4, NabFab, and anti-Fab Nb at 1.5 mg/ml in 20 mM Tris-HCl, pH 7.5, and 150 mM NaCl was prepared as described in the Methods. Images were collected using Titan Krios TEM with a K3 detector of S^2^C^2^, Stanford, CA. The particle reconstructions and modeling were performed as described in the methods. The final volume did not include the anti-Fab Nb during Local Refinement due to relatively poor densities. **(a)** The raw image after motion correction. **(b)** Representative 2D-Classes generated by CryoSPARC program. MelB_St_ in nanodiscs, Nb725_4, NabFab, and anti-Fab Nb can be easily recognized. **(c)** GSFSC resolution was calculated by cryoSPARC Validation (FSC) using two half maps generated by the CryoSPARC Local Refinement program. The number of particles used for the volume reconstruction was presented. **(d)** Particle distribution of orientations over azimuth and elevation angles generated by CryoSPARC Local Refinement program. **(e&f)** The structure of MelB_St_/Nb725_4/NaFab complex. The volume (*e*) and cartoon representation (*f*) were colored by polypeptide chains as indicated. Nanodiscs were transparent and colored in light gray. Sphere and sticks in the panel *f* highlighted Na^+^ and its ligands.

Various ligands binding to the Nb/MelB_St_ complex in the presence of Na^+^ were also examined with ITC. As published^29^, the melibiose binding was undetectable in the presence of Nb725 (**sFig. 3a**). To characterize the sugar effect quantitatively, the α-NPG binding was carried out. Interestingly, the exothermic binding observed in the absence of the Nb changed to endothermic binding reactions in the Nb-bound states; and also, the α-NPG affinity decreased by 21-or 32-fold for Nb725 or Nb725-4, respectively (**Table 2**; **sFig. 3b**). Inhibitions of galactosides have been observed in the case of EIIA^Glc^/MelB_St_ complex^33,34^. Collectively, galactosides binding to MelB_St_ were mutually exclusive with Nbs (Nb725, Nb725_4, or Nb733) or EIIA^Glc^, and the negative cooperativity implied that MelB_St_ conformations favored by galactosides and those binders differ. This conclusion was further supported by the Na^+^ or EIIA^Glc^ binding to the MelB_St_/Nb complexes (**Table 2**; **sFig. 3c-d)**, which showed no inhibition for Na^+^ affinity and even slightly better binding for EIIA^Glc^. Therefore, MelB_St_ complexed with Nb725_4 retains its physiological functions, and its conformation is expected to be similar when trapped by either of those Nbs and EIIA^Glc^ but is different from the sugar-bound outward-facing structure [PDB ID 7L17]^19^.

**Table 2.**
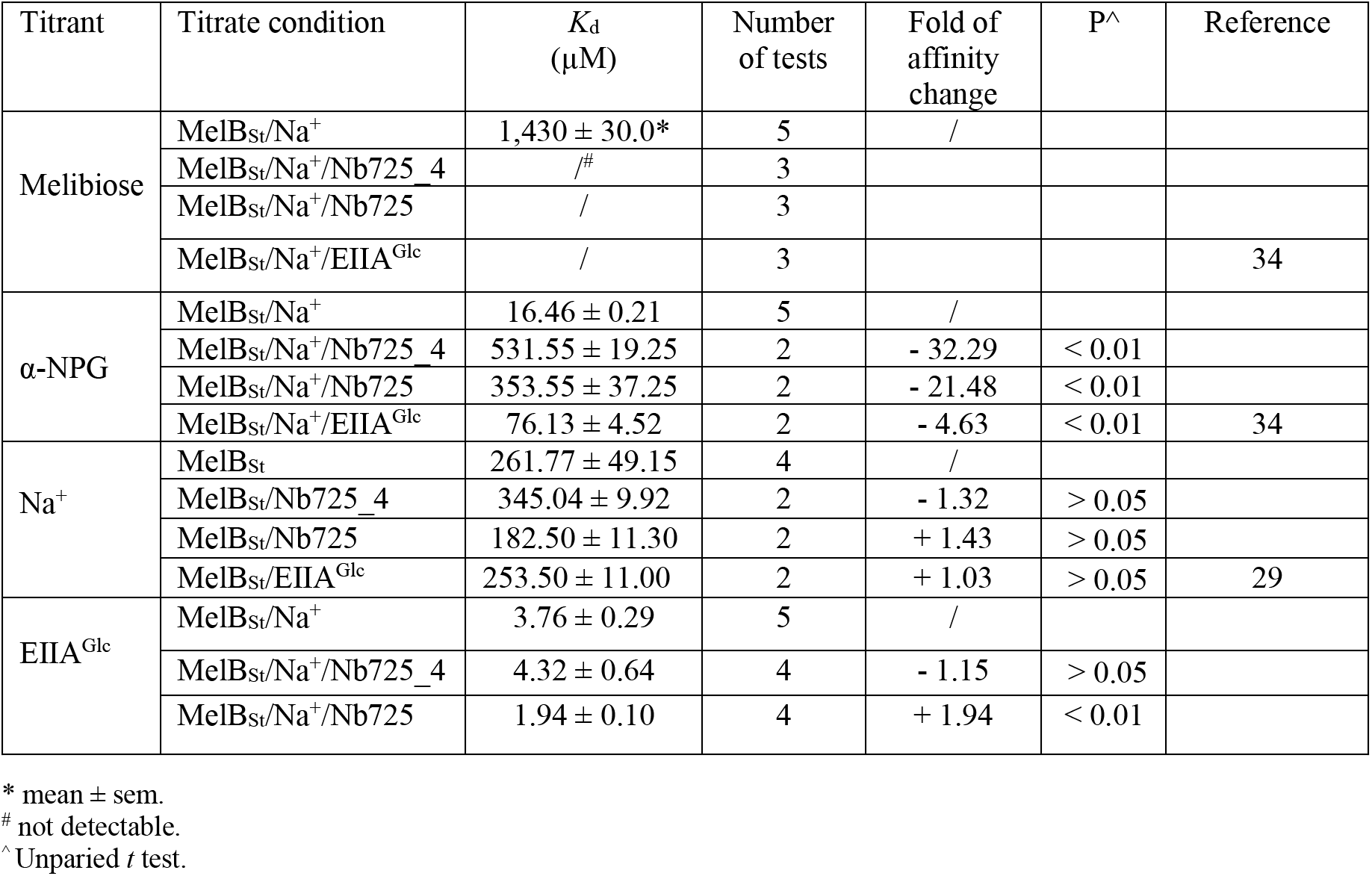
Nb effects on MelB_St_ binding to sugar, Na^+^ and EIIA^Glc^.

### Inward-facing MelB_St_ trapped by Nb725_4

CryoEM-SPA was used to image the complex consisting of MelB_St_ in nanodiscs, Nb725_4, NabFab, and anti-Fab Nb. The data collection, processing, and evaluation followed the standard protocols **(sFigs. 4-5)**. From a total number of 296,925 selected particles, structures of MelB_St_, Nb725_4, and NabFab were determined to a golden standard Fourier shell correlation (GSFSC) resolution of 3.29 Å at 0.143 (**Fig. 2**). The map for anti-Fab Nb were relatively poor and excluded from particle reconstruction. The quality of the Coulomb potential map allowed for unambiguous docking of available coordinates of NabFab [PDB ID 7PHP], a predicted model for Nb725_4 by Alpha-Fold 2^40,41^, and the N-terminal helix bundle from the outward-facing crystal structure of D59C MelB_St_ [PDB ID 7L17]. The C-terminal helix bundle of MelB_St_ and the ion Na^+^ were manually built. The final model contains 417 MelB_St_ residues (positions 2-210, 219-355, and 362-432), with 6 unassigned side-chains at the C-terminal domain (Leu293, Tyr355, Arg363, Tyr369, Tyr396, and Met410) due to the map disorder, 122 of Nb725_4 residues 2-123, and 229 of NabFab H-chain residues 1-214, and 210 of NabFab L-chain residues 4-213, respectively. The local resolution estimate showed that the cores of NabFab, Nb725_4, and the N-terminal helix bundle of MelB_St_ exhibit better resolutions up to 2.84 Å and most regions in the C-terminal helix bundle of MelB_St_ have resolutions between 3.2 to 3.6 Å (**sFig. 6a**). The N-terminal helix bundle has a completely connected density; however, the map corresponding to three cytoplasmic loops at positions 211-218 in the middle loop_6-7_, positions 356-361 in loop_10-11_, or the entire C-terminal tail after Tyr432, is missing. The map quality, model statistics, and the model-map matching *Q* scores, generally matched to this reporting resolution (**Supplementary sTable 1; sFig. 6a-c**).

The NabFab structure in this complex is virtually identical to that in the NorM complex [PDB ID 7PHP]^36^ with rmsd of all atoms of 1.334 Å (**sFig. 7a-c**). When the alignment focused on the NabFab H-chain, the two NabFab/Nb binary complexes in both complexes were organized similarly (**sFig. 7d)**, and the binding of Nbs with their corresponding transporters varied.

The structure shows that the Nb725_4 is bound to the cytoplasmic side of MelB_St_ at an inward-open conformation (**sFig. 8a&b)**, which supports the functional analyses and reveals the MelB_St_ conformation for the Nb binding. The contact surfaces match in shape and surface potential with a contact area of 435 A^2^. CDR-1 (Arg30, Asp31, Asn32, and Ala33) and CDR-2 (Tyr52, Asp53, Leu54, Tyr56, Thr57, and Ala58) form interactions with two cytoplasmic regions of MelB_St_, the tip between helices IV and V, i.e., loop_4-5_, (Pro132, Thr133, Asp137, Lys138, and Arg141) and the beginning region of the middle loop_6-7_ (Tyr205, Ser206, and Ser207) (**sFig. 8c**). The Nb binding was stabilized by salt-bridge and hydrogen-bonding interactions at both corners of this stretch of inter-protein contacts. In the published outward-open conformation of MelB_St_, such a binding surface for recognizing Nb725_4 does not exist. Most regions of the middle loop, the C-terminal helix bundle, and the C-terminal tail helix must be displaced to permit the binding, and some of those regions are completely disordered in this inward-facing structure. All data supported that Nb725_4 only binds to MelB_St_ in an inward-open state. Thus, it is a conformation-selective Nb.

Nb725_4, Nb725, and another Nb733^29^ behave similarly; all inhibit the sugar binding and transport. We conclude that all three binders are the inward-facing conformation-specific negative modulators of MelB_St_. Further, ITC measurements (**sFig. 2**) showed that MelB_St_ binds to either of three Nbs in the absence or presence of EIIA^Glc^ or binds to EIIA^Glc^ in the absence or presence of either Nbs. The complex composed of MelB_St_, EIIA^Glc^, and Nb725 (or Nb733^29^) can be isolated (**sFig. 9**).

### MelB_St_ trapped in a Na^+^-bound, low sugar-affinity state

Between helices-II&IV (**Fig. 3a&b**), one Na^+^ cation was modeled, liganded by four side-chains, Asp55, Asn58, Asp59 (helix-II), and Thr121 (helix-IV), at distances of between 2.1 - 2.8 Å. The backbone carbonyl oxygen of Asp55 is also in close proximity to the bound Na^+^. The local resolution of this binding pocket is approximately 2.9 - 3.2 Å. The model to map *Q*-scores for Na^+^ and the binding residues and side-chains are generally greater than the expected scores at this resolution except for the Thr121 side-chain, which has a lower score^42^. The observation of this cation-binding pocket is supported by previous extensive data^17,43,44^. Cys replacement at position Asp59 changes the symporter to a uniporter^19^. The D59C mutant does not bind to Na^+^ or Li^+^ as well as loses all three modes of cation-coupled transport but catalyzes melibiose fermentation in cells (melibiose concentration-driven transport)^19^ or melibiose exchange in membrane vesicles^11^. Asp55 also plays a critical role in the binding of Na^+^ or Li^+^ and in coupled transport^11,17^. Extensive studies of *E. coli* MelB also show that both negatively charged residues are involved in cation binding^21,45,46^. Thr121 has been determined to be critical for Na^+^, not H^+^ nor Li^+44^; and the Cys mutation at Asn58 also loses Na^+^ binding^11^. Interestingly, the Na^+^ recognition can be selectively eliminated by specific mutations. Further, the Na^+^ binding is supported by MD simulations at both inward and outward states using MD simulations in the absence of the Nb **(sFig. 10a-c)**, which reveals the ideal coordination geometry for Na^+^ binding.

**Fig. 3.**
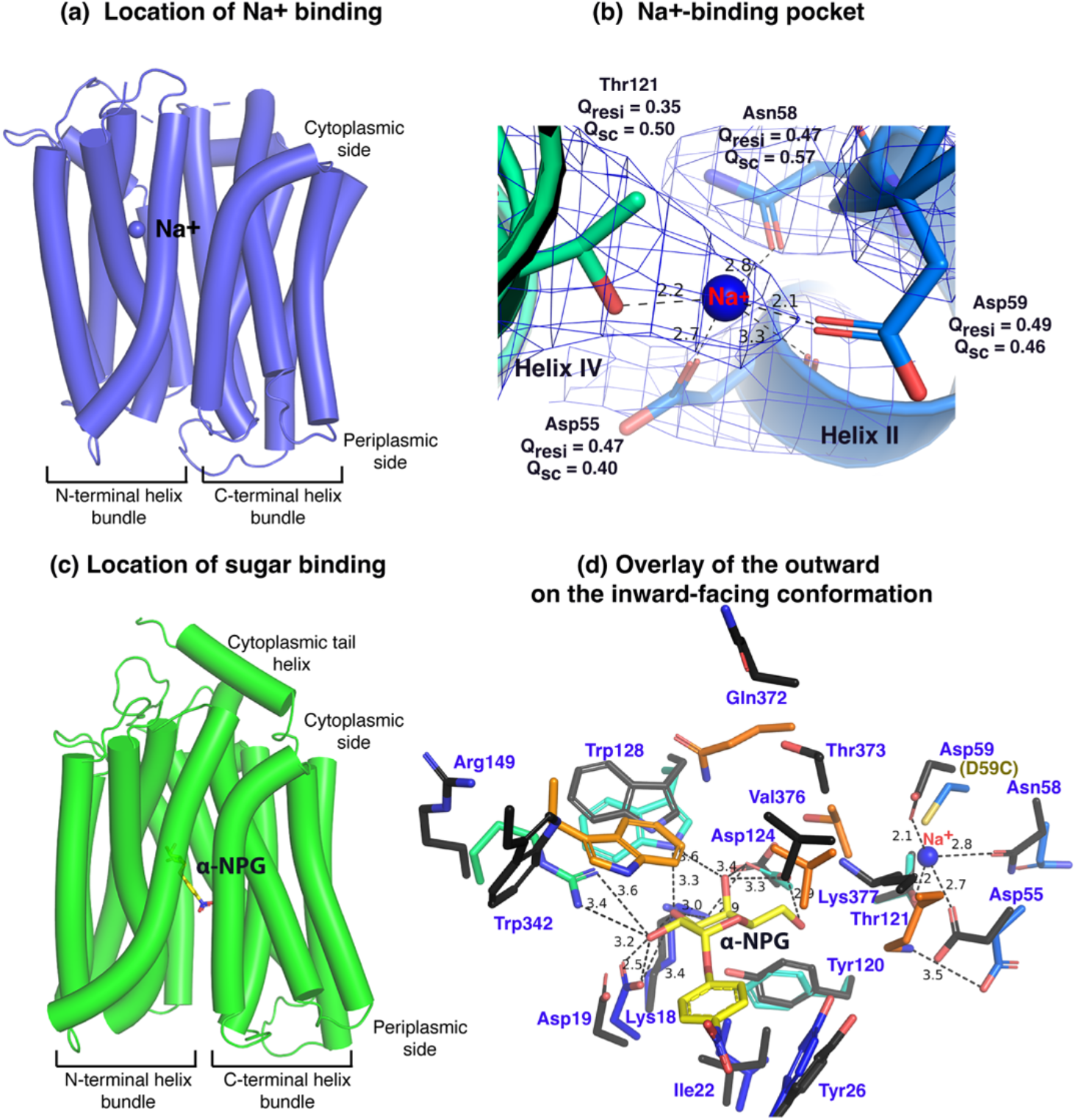
Na^+^-and sugar-binding pockets of MelB_St_. **(a)** Location of the Na^+^ binding site. The inward-facing cryoEM structure of the WT MelB_St_ was displayed in cylindrical helices with the cytoplasmic side on the top. One bound Na^+^ ion within the N-terminal helix bundle was shown in the blue sphere. **(b)** Na^+^-binding pocket. The isomesh map of the inward-facing conformation was created by the Pymol program using level 10 and carve of 1.8. The Na^+^ coordinates were shown in dash lines and interacting residues were shown in sticks. *Q*_res_, *Q* score for residue; *Q*_sc_, *Q* score for side chain. **(c)** Location of the galactoside-binding stie. The outward-facing x-ray crystal structure of D59C MelB_St_ mutant was displayed in cylindrical helices with the cytoplasmic side on the top [PDB ID 7L17]. One α-nitrophenyl galactoside (α-NPG) molecule was shown in the stick colored in yellow between the N-and C-terminal helix bundles. **(d)** Superimposed sugar-and cation-binding pockets. The a-NPG-bound outward-facing structure in *c* was aligned with the inward-facing cryoEM structure based on the 2-200 region. The residues in the sugar-and cation-binding pockets of the inward-facing cryoEM structure were colored in black and labeled in blue. D59C of the a-NPG-bound outward-facing structure is indicated in the parentheses. The a-NPG and Na^+^ were colored yellow and blue, respectively.

The crystal structure of sugar-bound D59C MelB_St_ (**Fig. 3c&d**; **sFig. 11)**, which retains an intact sugar-binding and translocation pathway, shows that the sugar-binding pocket is formed by both N-and C-terminal bundles^19^. Due to the Cys mutation, the cation-binding pocket is loosely packed and this mutant does not bind Na^+^ or Li^+^. When overlaying the outward-facing crystal structure upon the inward-facing cryoEM structures (**Fig. 3d**, **sticks in black**), little variation of the positioning of the sugar-binding residues on the N-terminal bundle is observed except for Arg149, but all C-terminal residues involved in the sugar-binding pocket show greater displacements. In the cation-site mutant, Lys377 forms a salt-bridge interaction with Asp55; in the Na^+^-bound structure, Lys377 is closer to Asp124, and Asp55 is liganded with Na^+^.

The loosely arranged sugar-binding residues observed in the Nb725_4-bound conformation provide the structural foundation for measured lower sugar affinities (**Table 2**), so this structure represents an intracellular Na^+^-bound sugar-releasing conformation. Since the positioning of the key residues for galactoside specificity (Lys18, Asp19, and Asp124) exhibits virtually no change, it is conceivable that this conformation retains the initial recognition of sugar for reversal reactions, a common feature for carrier transporters^25,47^.

### Conformational changes between inward-and outward-facing states

The outward-facing structure was trimmed to match the residues resolved in the inward structure. The superposition of the outward-and inward-facing MelB_St_ exhibited rmsd of all atoms of 5.035 Å (**sFig. 12a)**, and the major displacement exists at both ends of transmembrane helices. The alignment based on the N-terminal positions 2-200 or the C-terminal positions 231-432 exhibits a clear difference in the angle of the domain packing (**sFig. 12b&c)**. The focused alignment of positions 2-200 or 231-432 exhibits greatly reduced rmsd values (**sFig. 12d&e**). The N-terminal domains at both conformations are virtually the same, especially for the helices-I, -II, and -IV that host the sugar-and cation-specificity determinant pockets (**sFig. 12f**). Some structural rearrangements within 3-5 Å exist in the C-terminal bundle, especially on both ends of cavity-lining helices (VII, VIII, X, and XI) and their link or extended loop (**sFig. 12g&h**). Thus, the conformational changes between the outward-and inward-facing states can be adequately described by a rocker-switch-like movement with some structural rearrangements in the C-terminal bundle as also indicated in **sFig. 13**.

### Substrate translocation barriers

The two structures clearly reveal the two essential barriers critical for substrate translocation, the thicker inner barrier in the outward structure and the thinner outer barrier in the inward structure (**Fig. 4a&d**). The core of the inner barrier is formed by three-helix pairs from both domains (V/VIII, IV/X, and back II/XI), and stabilized by an extensive charged network (**Fig. 4b**)^11,16,19,48^. The N-terminal Arg141 (helix-V) forms salt-bridge interactions with the C-terminal Asp351 and Asp354 (helix-X), and this core interaction links with four other charged residues to stabilize the cytoplasmic closure by forming an inner barrier. In addition, the inner barrier is also stabilized by two peripheral amphiphilic helices at the cytoplasmic side (in the middle loop and the C-terminal tail) (**Fig. 4b**). The entire middle loop_6-7_ across the two domains interact with broader areas (the N-terminal loop_2-3_, loop_4-5_, and helices-II & -IV, as well as the C-terminal loop_10-11_, helices-X & - XI, and the C-terminal tail). The C-terminal tail helix interacts with both N-and C-terminal domains (helices-V &-X). The contacting area between the two bundles (based on positions 2-212 and 213-450) is 2218 Å^2^ with nine salt-bridge and five hydrogen-bounding interactions.

**Fig. 4.**
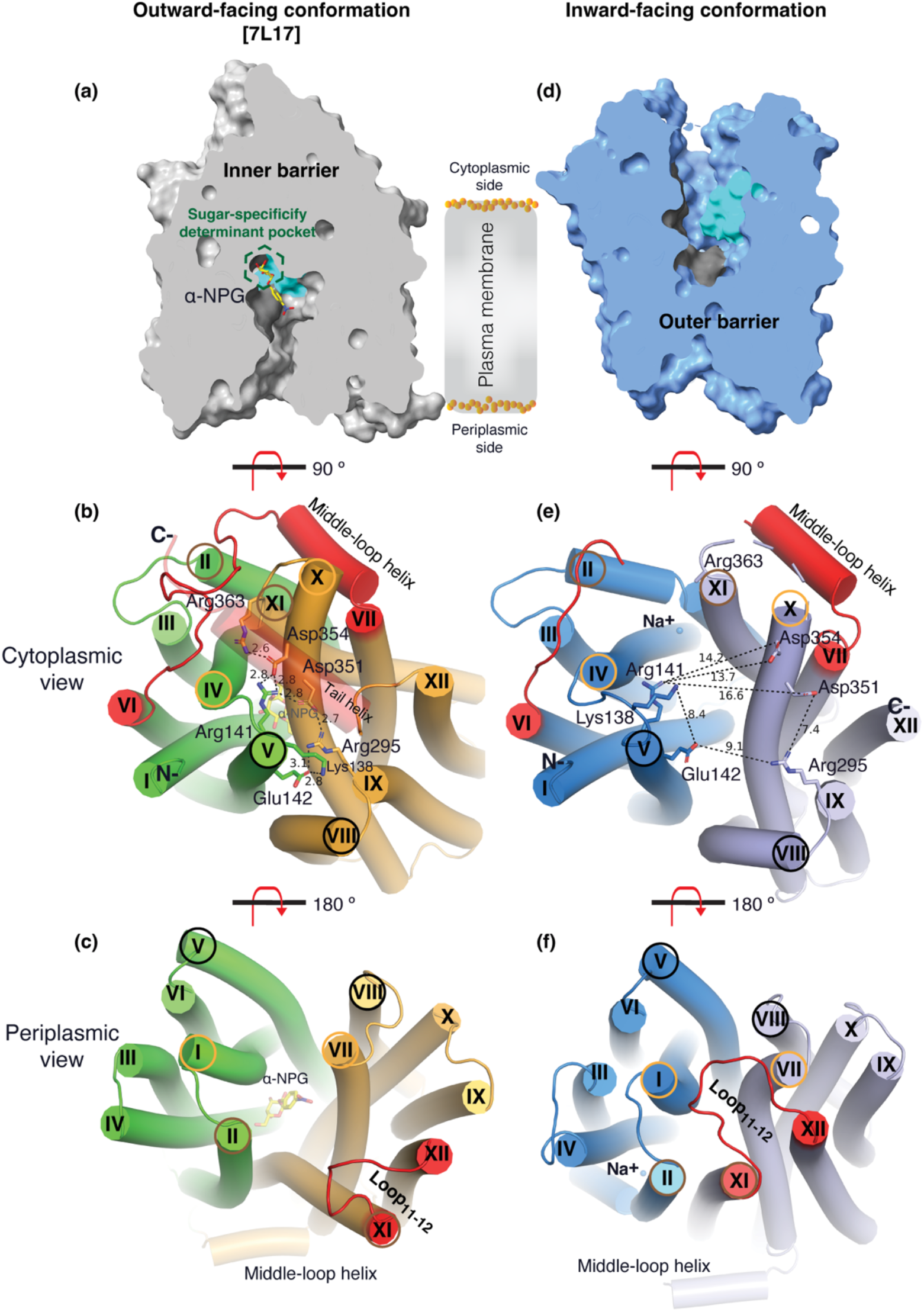
Barriers and sugar-binding pocket. Outward-facing [PDB ID 7L17; left column] and inward-facing [right column] structures were used to prepare the figures. **(a&d)** Side view with cytoplasmic side on top. The inner and outer barriers were labeled. The sugar-specificity determinant pocket was highlighted in a dashed hexagon. The residues contributing to the bound a-NPG in N-and C-terminal bundles were colored in dark gray and cyan, respectively. **(b&e)** Cytoplasmic view. The charged network between the N-and C-terminal bundles were colored in green and bright orange in panel *b*, or blue and light blue in panel *e*, respectively. The C-terminal tail helix was set in transparent in panel *b* but disordered in the inward-facing conformation in panel *e*. The charged residues were highlighted in sticks. Arg363 side chain missed the sidechain in the inward-facing structure. **(c&f)** Periplasmic view. The paired helices involved in either barrier formation were highlighted in the same colored circles. The α-NPG was colored in yellow and Na^+^ was shown in blue sphere. The cytoplasmic middle loop, the C-terminal tail, and the periplasmic loop_11-12_ were highlighted in red.

The core of the outer barrier in the inward-facing structure is also formed by three pairs of helices from both domains (V/VIII, I/VII, and II/XI) (**Fig. 4f**). The longest loop_11-12_ at the periplasmic side moves into the middle area and forms contacts with helices-I and -VII, sealing the periplasmic opening. Notably, helical pair I/VII or IV/X is only involved in the outer or inner barrier, respectively, and the other two cavity-lining helical pairs (V/VIII and II/XI) are engaged in both barriers (**Fig. 4b, c, & e, f**). The contact area between the two bundles at the periplasmic side of 1435 Å^2^ is much smaller. These structural features further supported that the outward-facing state is more stable than the inward-facing MelB_St_. In the inward-facing conformation, all of the salt-bridge interactions are broken (**Fig. 4b&e**), and the Arg141 and Lys138 engage in the binding of Nb725_4 (**sFig. 8c**). Therefore, this cytoplasmic-side salt-bridge network is the inner barrier-specific.

Assessed by the sugar-bound structure (**Fig. 4a**), the substrate-binding pocket is outlined by Trp128 and Trp 342 on the inner barrier and Tyr26 on the outer barrier (**Fig. 3d**; **sFigs. 11, 12f**), part of both barriers. Notably, the major binding residues that involve the multiple hydrogen-bonding interactions with the galactosyl moiety are only part of the inner barrier. In the inward-facing state, the inner barrier is broken, which exposes the binding site to the cytoplasm and also results in the interruption of the sugar-binding pocket (**Fig. 4d**). The C-terminal binding residues moved away due to the displacement of the backbone of helices-X/XI, and the sugar-binding residues become loosely organized with a low sugar-binding affinity. The structural features indicated that the molecular mechanisms for the sugar translocation simply result from the barrier switch generating a low sugar-affinity inward state of MelB. The outer barrier does not contain residues essential for galactoside binding, so the break of the outer barrier has a much less profound effect on the sugar-binding affinity.

### Conformational dynamics measured by HDX-MS

Bottom-up HDX-MS measures the exchange rate of amide hydrogens with deuterium by determination of time-dependent peptide level mass spectra^37^. It is a label-free quantitative technique that can disclose the dynamics of a full-length protein simultaneously. This approach was utilized to determine MelB_St_ dynamics in this study. The Na^+^-bound WT MelB_St_ proteins exist in multiple conformations including at least the outward-and inward-facing states but strongly favor the outward-facing states. The Nb725_4-trapped MelB_St_ exists only in an inward-facing conformation. A pair of WT MelB_St_ and WT MelB_St_ complexed with Nb725_4 was subjected to in-solution HDX-MS experiments. MelB_St_ yields 86% coverage with 155 overlapping peptides at an average peptide length of 9.3 residues (**Fig. 5a&b**; **sTable 2**), leaving 66 non-covered residues; most are located in the middle regions of transmembrane helices or in a few cytoplasmic loops (**Fig. 6**, **ball in gray**).

**Fig. 5.**
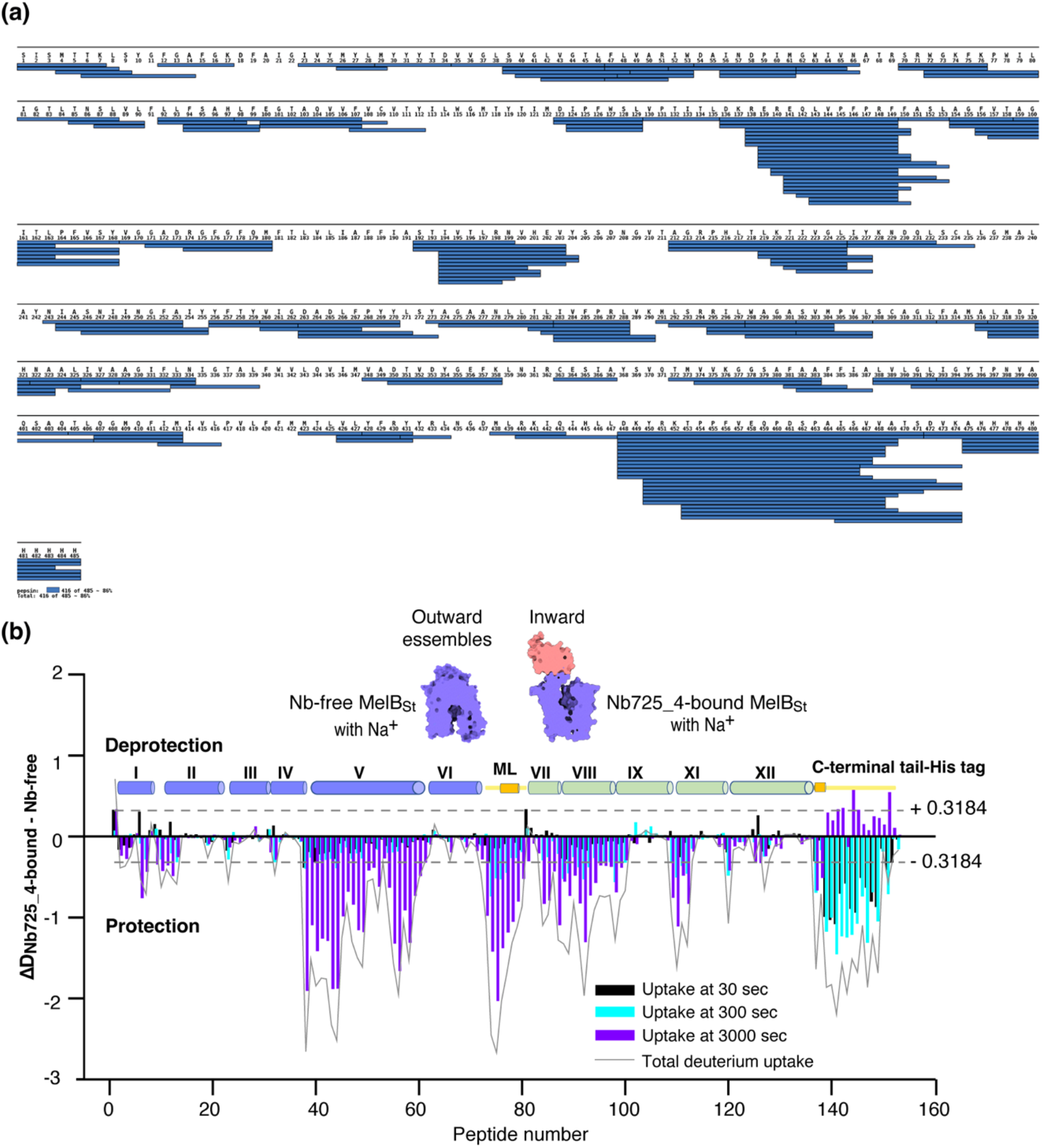
MelB_St_ dynamics probed by HDX-MS. **(a)** MelB_St_ peptide sequence coverage. The peptides of the deuterated MelB_St_ were determined based on the MelB_St_ peptide database that was generated by nonspecific digestions of non-deuterated MelB_St_ as described in the Methods. Peptides were confirmed in the HDX-MS experiment. Blue bar, the covering of each peptide. The amino-acid sequencing identification number should be -1 for each position due to the processed Met at position 1. The 10xHis Tag was included in the data analysis. **(b)** Residual plots (D_Nb725_4-bound - Nb-free_) against the overlapping peptide numbers for each time point and the sum of uptake. MelB_St_ alone or bound with Nb725_4 in the presence of Na^+^ were used to carry out the HDX reactions as described in the Methods. Black, cyan, and blue bars, the deuterium uptake at 30, 300, 3000 sec, respectively; gray curve, the sum of uptake from all three time points. Deprotection, ΔD (D_Nb725_4-bound – Nb-free_) > 0; protection, ΔD < 0. Each sample was analyzed in triplicates. Cylinders indicated the helices; the transmembrane helices were labeled in Rome numeral. The length of the cylinder does not reflect the length of corresponding helices but is estimated for mapping the deuterium-labeled overlapping peptides. The uncovered regions were not included. ML (cytoplasmic middle loop) and C-terminal tail including the His tag were colored in yellow. Dashed lines, the threshold.

**Fig. 6.**
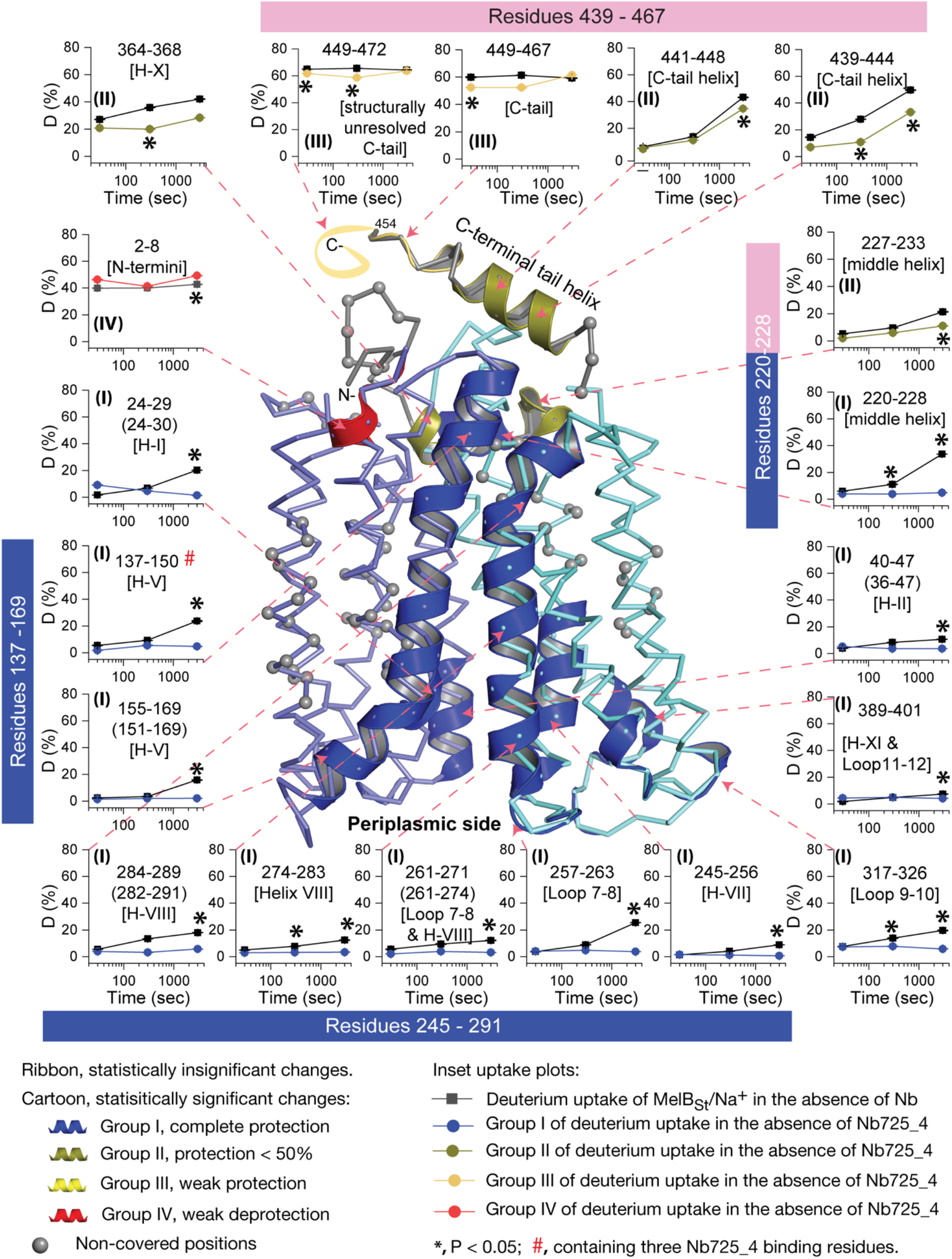
Peptide mapping of HDX results. HDX results were presented in Fig. 5. Any peptide of ΔD with P ≤ 0.05 and ∆D ≥ |0.3184| were treated as significant. The peptides with statistically significant differences at the 3000 sec time point were mapped on the outward-facing structure [PDB ID 7L17]. **Inset,** the deuterium uptake time course of representative peptides was plotted as a percentage of deuterium uptake relative to the theoretic maximum number (D%). The peptides, either as a single peptide or a group of overlapping peptides (in the parentheses), are labeled and indicated in the structure by the pink arrow in dashed lines. In the brackets, the corresponding secondary structure or loops. Error bar, sem; the number of tests, three. Other symbols were presented within the figure.

The effect of MelB_St_ structural dynamics imposed by Nb725_4 is presented as the differential deuterium labeling (ΔD). As shown in the residual plot of each peptide between the two states (D_Nb725_4-bound – Nb-free_) (**Fig. 5b**) at three labeling time points, variable levels of protections (less deuterium exchange in the complex state) are observed in MelB_St_. A slight deprotection is only observed at the N-terminus. After proper statistical analysis, there are 232 positions (>50% labeled positions) exhibiting insignificant HDX changes (D_Nb725_4-bound – Nb-free_ value < |0.3184| or P > 0.05) as indicated by ribbon representation (**Fig. 5b**). The remaining 177 positions, not counting the His tag, covered by 71 overlapping peptides show statistically significant changes (D_Nb725_4-bound – Nb-free_ value ≥ |0.3184| and P ≤ 0.05) and are highlighted by the cartoon representation (**Fig. 6**). Overall, those regions are mainly distributed on the cavity-lining helices and cytoplasmic peripheral regions including the middle loop and C-terminal tails, as well as the C-terminal periplasmic loops. The corresponding deuterium uptake curves covering 3-magnitude time points were plotted **(Extended Fig. 1)** and the representative peptides were displayed as insets around the structure (**Fig. 6**, **insets)**. Most Nb725_4 binding residues are not covered, either not covered by the labeled peptides (Tyr205, Ser206, and Ser207) or by insignificant change (Pro132 and Thr133), except for Asp137, Lys138, and Arg141 that were presented in 8 peptides. Depending upon the exchange rate and response to the binding of Nb725_4, the MelB_St_ HDX can be categorized into four groups.

<u>Group-I</u> peptides show low-level HDX exchange rates and Nb725_4 afforded complete inhibition for up to at least 50 min (**Fig. 6**, **cartoon and curve in blue**). These peptides are located in transmembrane helices, middle-loop helix, and C-terminal periplasmic loops across both domains and most are clustered at the cavity-lining helices. Peptides carrying the Nb-binding residues Asp137, Lys138, and Arg141 are marked in the plots (**Fig. 6**, **inset; Extended Fig. 1**). <u>Group II</u> peptides show faster deuterium uptake over time (**Fig. 6**, **cartoon and curve in deep olive**). The Nb725_4 binding affords small to moderate protections. The group-II peptides are spotted at three cytoplasmic regions; interestingly, all surround the helix-X, which plays a critical role in the stability of the inner barrier. At the inward-facing structure, most of those residues are either missing or poorly resolved. The deuterium uptakes in the Nb725_4-bound MelB_St_ complex likely result from structural flexibility.

<u>Group-III</u> peptides show higher-level HDX exchange rates of greater than 60% for both bound and free states even at 30 sec (**Fig. 6**, **drawing in yellow**). Nb725_4 only affords short-time, weak protections. They are located in the C-terminal tail unstructured region, and most of them are structurally unresolved due to severe disorders. <u>Group-VI</u> peptide only contains the short N-terminal tail showing a medium-level uptake (40-50%) across all labeling times, and Nb725_4 affords a slight deprotection due to allosteric changes upon the Nb725_4 binding (**Fig. 6**, **cartoon and curve in red**). The results show that Group III and IV peptides covering both tails are solvent-accessible dynamic regions, which is consistent with the notion that they play little role in transport-critical conformational transitions.

The 30-sec deuterium uptake in the absence of Nb725_4 for the first two groups was further analyzed. Among the five group-I and four group-II members with faster exchanges as judged by the shortest reaction time of 30 sec (**sFig. 14)**, except for the two periplasmic loop_7-8_ (peptide 261-271) and loop_9-10_ (peptide 317-326), others are clustered on the cytoplasmic side. In the outward-facing structure (**Fig. 7a**), the spotted peptides pack against one another: helix-V (peptide 137-150) with helix-VIII (peptide 284-289), and the middle-loop helix (peptide 220-233) with helix-X (peptide 364-368). All surround the conformation-dependent charged network. Notably, this short peptide 137-150 links the sugar-binding residue Arg149 to the charged network by residues Lys138, Arg141, and Glu142 (**Fig. 4b**). In the inward-facing structure (**Fig. 7b**), these packed helices were moved against one another. Therefore, these higher dynamic regions may play important roles in responding to the ligand binding and initiating the global conformational transition.

**Fig. 7.**
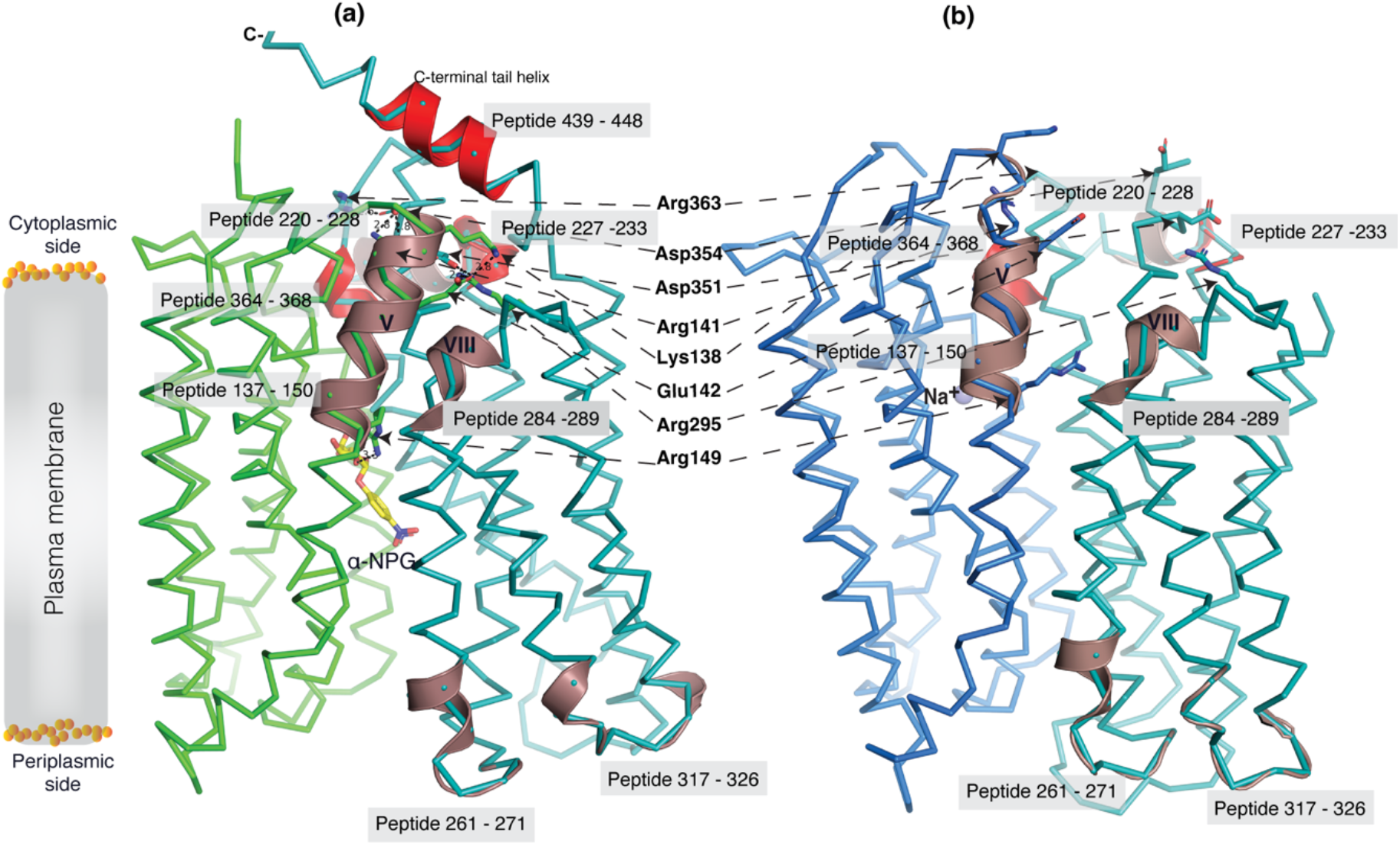
Dynamic regions of MelB_St_. The peptides that exhibited faster HDX rates (> 5% at 30 sec) were mapped on both the a-NPG-bound outward-facing structure [7L17] in panel ***a*** and the inward-facing cryoEM Nb725_4-bound structure in the panel ***b***, respectively. All peptides were labeled and highlighted in a transparent gray box. The charged residues forming the inner barrier-specific salt-bridge network were highlighted in sticks and indicated by black arrows in dashed lines. In the Nb725_4-bound structure (***b***), the C-terminal tail was disordered. The cartoon colored in dirtyviolet, the group I peptides with faster HDX rate; the cartoon colored in red, group II peptides. The α-NPG was highlighted in yellow and Na^+^ was shown in blue sphere.

## Discussion

Formation and stabilization of a high-energy inward-facing conformation of MelB_St_ in a form suitable for structure determination by cryoEM-SPA was a technically challenging sample preparation problem, which we solved by the formation of a complex of MelB_St_ with a conformation-selective binder and fiducial NabFab. The resulting structure, a high-energy inward-facing, low sugar-affinity state of MelB_St_ with a bound Na^+^, represents the intracellular sugar-releasing/rebinding state in the stepped-binding model for the mechanism of coupled transport (**Fig. 8**)^15,19^. Our previous results on the cooperativity of binding^28^ supported the contention that Na^+^ binding increases the melibiose affinity to allow melibiose uptake at lower available concentrations, a critical evolved mechanism for cation-coupled transporters. The new Na^+^-bound sugar-releasing inward-facing cryoEM structure provides a structural basis to support the transport model based on a sequential process of the sugar-binding after Na^+^ binding first and sugar release prior to Na^+^ release (**Fig. 8**).

**Fig. 8.**
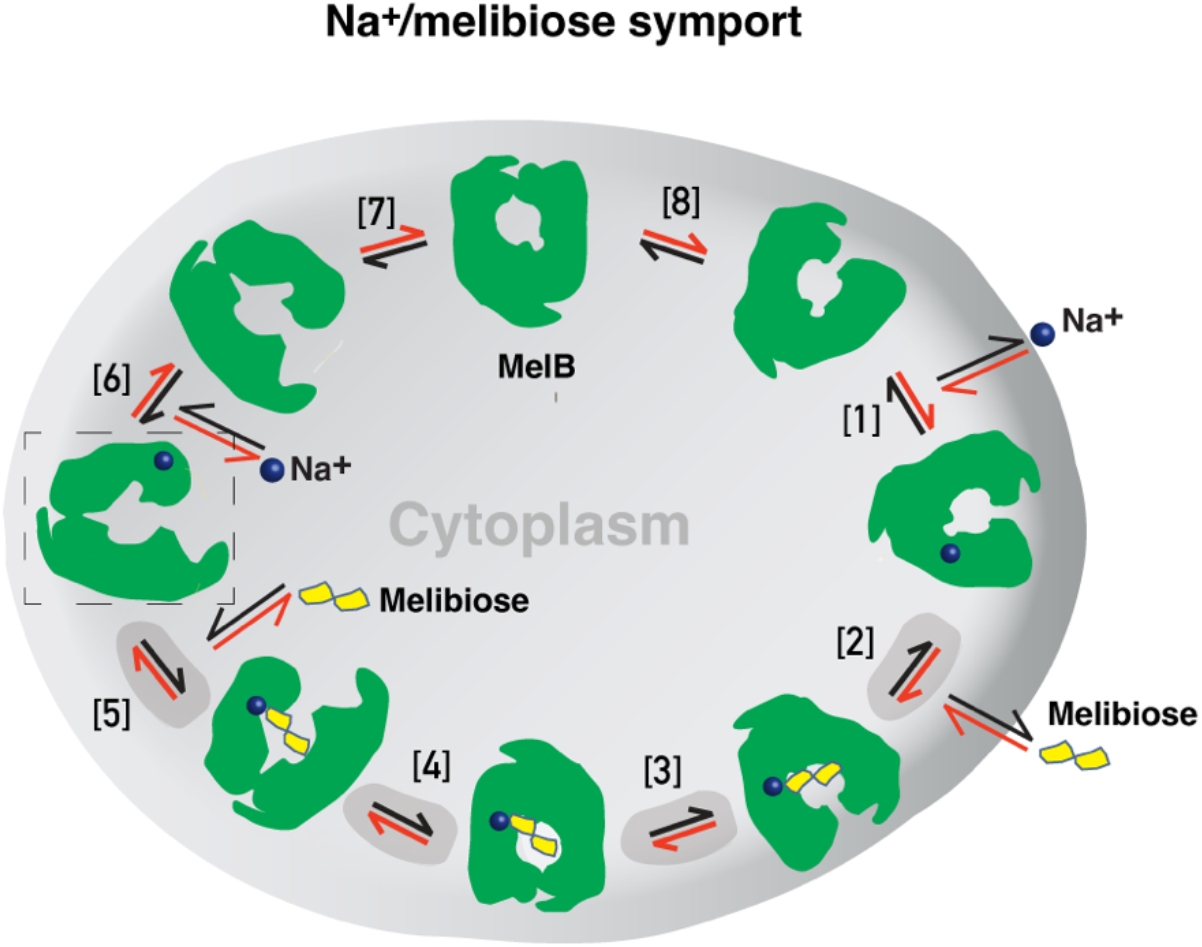
Stepped-binding model for the Na^+^/melibiose symport catalyzed by MelB. Eight states are postulated including transient intermediates. In this reversal reaction, the cation binds prior to the sugar and releases after sugar release. Melibiose active transport or inflex begins at step [1] and proceeds via the red arrows clockwise around the circle, with one melibiose and one cation inwardly across the membrane per cycle. Melibiose efflux begins at step [6] and proceeds via the black arrows anti-clockwise around the circle, with one melibiose and one cation outwardly across the membrane per circle. Melibiose exchange begins at step 6, and only takes 4 steps involving steps [2 – 5] as highlighted in gray color. The low-sugar affinity inward-facing Na^+^-bound cryoEM structure represents the state after the release of sugar, as indicated by a dashed box.

The alternating-access movement has been widely used to describe the conformational change of transporters^3,6,7,12,13,26,27^. This description focuses on how the substrates access their binding site or leave the protein. The core symport mechanism for coupled substrate translocations is the mobile barriers involving the inner and outer barriers as proposed originally by Peter Mitchell^49^ (**Fig. 4a&d**). Our binding thermodynamic cycle study on the determination of heat capacity change (∆*C*_p_)^28^ showed positive values with the binding of one substrate, suggesting the dominance of hydrations and opening of the transporter, and negative values with the second substrate binding, suggesting dehydration and transition to the occluded intermediate state as indicated at step [3] in Figure 8. The data fit well with a barrier mechanism that prevents a single substrate molecule from crossing the protein to favor obligatory co-translocation. When this inner barrier breaks as a result of the conformational switch induced by the binding of two substrates, the sugar-binding specific pocket which is part of this inner barrier also breaks (**Fig. 4e**), creating the intracellular sugar low-affinity state and the sugar-releasing path. Collectively, based on structural and thermodynamical analyses, all show that cooperative binding is integral to barrier switching in the core symport mechanism.

For substrate translocation, historically, a two-binding-site walking model was proposed: a high-affinity site at the outer surface and a low-affinity site at the inner surface. In this functional and structural paradigm, a substrate molecule moves across the protein from its high-affinity binding site to a low-affinity site for subsequent release. This canonical model has been challenged in LacY as the same binding affinity was measured at both the inner and outer surfaces^50^, and a single substrate-binding site was also observed in its 3D structures^3,6,26^. Therefore, a single-binding site in the middle of the protein has been the current canonical model. Actually, one of the major challenges in measuring the substrate binding in membrane transporters is the intrinsic dynamic feature of the transport proteins, which makes it difficult to attribute the outcome to a specific state. Therefore, the substrate binding affinity measured in a bulk sample is an average of binding affinities from all of the individual molecules in a distribution. It is also challenging to establish a correlation between measured binding affinities and experimentally determined structures. We approached this problem by using the conformation-specific binding protein Nb725_4 to decouple the binding and conformational dynamics shifting conformers to a single state or a narrowed cluster, i.e., through experimental modulation of the distribution. The results of the structural determination and binding analysis in the combination of the conformation-selective Nb permitted the extension of the current single substrate-binding site (and single affinity) model to a more refined model of one site possessing conformation-dependent higher-affinity and lower-affinity states. Identification of this critical lower-affinity inward-open state of the sugar-binding site is significant for understanding the molecular basis of higher intracellular substrate accumulation.

The shared cation site for Na^+^, Li^+^, or H^+^, which is hosted only by the N-terminal two helices (II and IV), is in close proximity but not part of the mobile barrier at either outward-and inward-facing state. The inward-facing state retains the Na^+^ binding affinity (**Fig. 1d**; **Table 2**) and the bound Na^+^ ion was observed (**Fig. 3ab**). Consistently, the simulated Na^+^-binding site at both outward-and inward-facing conformations is virtually identical (**sFig. 10**), which is in stark contrast to the conformation-dependent sugar binding in MelB_St_. No conformational dependency of Na^+^ binding might suggest the mechanism for the reversible reaction. It is likely that this Na^+^-binding site co-evolved with the *E. coli* intracellular Na^+^ homeostasis maintained to 3-5 mM^51^ to achieve optimal activity.

To interrogate the structural dynamics of MelB_St_ and conformational switch mechanisms, an HDX-MS comparability study in the absence or presence of the conformational Nb725_4 was performed. HDX-MS is a powerful approach for characterizing protein conformational flexibility and dynamics^37,38^ and has recently been applied to membrane transport proteins^52–54^. Under experimental conditions, two major parameters (solvent accessibility and hydrogen protonation) contribute to the H/D exchange rate. The residual plot clearly indicates that the major effects of Nb725_4 binding are the reduction of the conformational dynamics of MelB_St_ (**Fig. 5b**). Stronger protections are observed at both transmembrane helix bundles, especially both ends of the cavity-lining helices (**Fig. 6**, **cartoon in blue**), which marks the most conformationally dynamic regions that correlate to the conformational transition in MelB_St_. Notably, the Nb-free Na^+^-bound WT MelB_St_ exists in both outward-and inward-facing conformations with a bias towards the outward-facing state^11,19^, and the MelB_St_ in the Nb725_4 complex exists only as an inward-facing conformational ensemble. Therefore, the differences observed by HDX-MS between the two samples in the absence and presence of Nb725_4 likely disclosed, to some extent, the MelB_St_ global conformational transitions, in addition to the atomic motions.

While the resolution of HDX is at a peptide level, the data showed marked effects on two well-conserved cytoplasmic helices of MFS transporters (on the middle loop and the C-terminal tail). Those peripheral amphiphilic helices are likely to play important roles in the stability of the inner barrier. Consistently, a recent atomic force microscopy study has shown that the middle loop becomes loosely packed in the presence of both sugar and cation^20^.

As described, the peptide 137-150 of helix-V contains a sugar-binding residue Arg149 and three conformation-critical residues Lys138, Arg141, and Glu142 in the inner barrier-specific charged network (**Fig. 4**). Extensive site-directed mutagenesis and second-site suppression showed that this charged network play important role in the conformational changes and the mutations at all three Arg at positions 141, 295 and 363 (**Fig. 4b**) can be substantially rescued by a single mutation D35E (helix-I) located at the other side of the membrane^48^. Arg141 and Glu142 of *E. coli* MelB have been shown to participate in the conformational transition^55^. Pro148 has been identified by second-site mutation to reduce the transport *V*_max_ of a *V*_max_ elevated mutant by modulating the rate of conformational change^56^. Thus, the helix-V dynamic region (peptide 137-150) could link the sugar-binding pocket with the inner barrier-specific salt-bridge network. Together with the interactions with the dynamic regions (residues 284-289) of helix-VIII, the substrate binding occupancy could be propagated, via an allosteric mechanism, from the sugar-binding pocket to the cytoplasmic peripheral helices as a pre-transition for the opening of the transporter to the cytoplasm^20^.

All of the conformation-selective nanobodies (Nb725, Nb733, and the hybrid Nb725_4) are expected to bind MelB_St_ similarly and trap it at the high-energy state, which is consistent with the observation of very low-frequency occurrence during Nbs selection^29^. The effects of the Nbs binding mimic the regulatory effect imposed by EIIA^Glc^ binding^33,34^, i.e., reduction of sugar-binding affinity with no change in the Na^+^ binding. Interestingly, either Nbs or EIIA^Glc^ can bind to MelB_St_ in the absence or presence of the other (**Table 2**; **sFigs. 2&3**); the complex composed of all three proteins can be isolated (**sFig. 9**)^29^. The extensive comparisons strongly support the notion that the Nbs and EIIA^Glc^ can concurrently recognize two spatially distinct epitopes of MelB_St_, and the inward-facing conformation trapped by the Nb mimics the conformation of MelB_St_ under the EIIA^Glc^ regulation. Available data showed that three positions (Asp438, Arg441, and Ile445) on one face of the C-terminal tail helix of MelB_St_ might be involved in EIIA^Glc^ binding^57^, and this region is conserved with the later known EIIA^Glc^ binding site in MalK^35^. Thus, the cytoplasmic surface of the C-terminal domain could potentially support the EIIA^Glc^ binding at the inward-facing state; unfortunately, the C-terminal tail helix is disordered in this Nb-bound inward-facing structure,

In this study, we demonstrated that the mobile barrier mechanism plays a central role in substate translocation, the barrier dynamics are modulated by the cooperative binding of substrates, and the inner/outer barrier shifting directly regulates the sugar-binding affinity, with no effect on the Na^+^ binding. Furthermore, we proposed that the functional inhibition of this MFS transporter by the central regulatory protein EIIA^Glc^ is achieved by trapping MelB_St_ in a low-sugar affinity inward-facing state. Overall, our studies provide substantial information to explain the obligatorily coupled transport of Na^+^ and sugar substrate in structural and dynamic terms collectively.

## Methods

### Reagents

MacConkey agar media (lactose-free) was purchased from *Difco.* Unlabeled melibiose and 4-nitrophenyl-α-D-galactopyranoside (α-NPG) were purchased from Sigma-Aldrich. Maltose was purchased from *Acros Organics* (*Fisher Scientific*). [1-^3^H]Melibiose (5.32 Ci/mmol) was custom synthesized by *PerkinElmer.* Detergents undecyl-β-D-maltopyranoside (UDM) and n-dodecyl-β-D-maltopyranoside (DDM) were purchased from *Anatrace*. *E. coli* lipids (Extract Polar, 100600) were purchased from *Avanti Polar Lipids, Inc*. Oligodeoxynucleotides were synthesized by *Integrated DNA Technologies*. LC-MS grade water, LC-MS 0.1% formic, LC-MS grade acetonitrile with 0.1% formic acid in water were purchased from Fisher Scientific (Hampton, NH). Guanidine hydrochloride, citric acid, and Zirconium (IV) oxide 5-μm powder were purchased from *Sigma-Aldrich* (St. Louis, MO). Deuterium oxide (99^+^% D) was purchased from Cambridge Isotope Laboratories (Tewksbury, MA). All other materials were reagent grade and obtained from commercial sources.

### Strains, plasmids, and primers

The genotype and source of *E. coli* strains used in this study are described in Supplementary **sTable 3** unless otherwise described specifically. The commercial *E. coli Stellar^TM^ or XL1 Blue* were used for plasmid construction. The *E. coli* DB 3.1 strain was used to construct the *ccdB*-containing universal FX Cloning vectors^58^. *E. coli* DW2 *(melB^-^lacY^-^*)^59^ was used for functional studies, and DH5*α cyaA*^-^ strain was used for *in vivo* protein-protein interaction assay^29^. *E. coli* BL21 (DE3) strain and *E. coli* ArcticExpress (DE3) strain were used for overexpression nanobodies; *E. coli* BL21 (DE3) C43 strain^60^ was used for overexpression NabFab; *E. coli* BL21(DE3) T7 express strain was used to express EIIA^Glc33^. The plasmids used or created and primers in this study were also listed in **sTable 3**.

### CDR grafting

CDR grafting technique^39^ to transfer the binding specificity of MelB_St_ Nbs to the TC-Nb4, which can be recognized by the NabFab, was performed^36^ (**sFig. 1a**). The DNA fragment, which carried the hybrid Nb725_4 along with N-terminal pelB signal peptide (MKYLLPTAAAGLLLLAAQPAMA) and C-terminal HRV-3C protease site (LEVLFQGP) and 8xHis-tag, was synthesized, digested using restriction enzymes NdeI and XhaI, and ligated with the corresponding sites on expression plasmid pET26b+, resulting in the pET26/Nb725_4.

### MelB_St_ protein expression and purification

A 10-L fermenter was used for the constitutive expression of MelB_St_ encoded by the expression plasmid pK95ΔAH/MelB_St_/CHis_10_ in the *E. coli* DW2 cells using Luria-Bertani (LB) broth supplemented with 50 mM KPi, pH 7.0, 45 mM (NH_4_)SO_4_, 0.5% glycerol, and 100 mg/L ampicillin. The cultures were grown at 30 °C until *A*_280_ ∼5. The procedures for membrane preparation, extraction with 1.5% UDM or 2% DDM, and cobalt-affinity purifications were performed as previously described^11,17,29^. The eluted MelB_St_ proteins were dialyzed overnight against a buffer of 20 mM Tris-HCl, pH 7.5, 100 mM NaCl, 10% glycerol, and 0.35% UDM or 0.01% DDM, and concentrated with Vivaspin 50 kDa MWCO membrane to approximately 50 mg/mL, stored at -80 °C after flash-frozen with liquid nitrogen.

### Overexpression and purification of membrane scaffold protein 1E3D1 (MSP1E3D1)

An expression plasmid pMSP1E3D1 (*Addgene;* plasmid 20066) was used for the overexpression of the MSP1D1E3 with a 7-His-tag and a TEV protease site in the *E. coli* BL21 (DE3)^61,62^. The culture was grown in LB media containing 0.5% glucose at 37 °C, induced by adding 1 mM IPTG at *A*_600_ of about 0.6, and incubated for another 2.5 hrs. The MSP proteins from the cell lysate were purified with INDIGO Ni-Agarose (Cube biotech) using a protocol as described^28,63^. The eluted proteins were dialyzed against 20 mM Tris-HCl, pH 7.5, and 100 mM NaCl and concentrated to ∼ 8 mg/ ml. The His-tag of MSP1E3D1 was removed by a His-tagged TEV protease (at 1:20 mole/mole ratio of TEV:MSP1E3D1) in the same buffer. The His-tagged TEV protease, un-processed His-tagged MSP1E3D1, and free His tag fragments were trapped within the Ni-Agarose column. The processed MSP1E3D1 proteins without a His tag in the flow-through were concentrated to about 6-8 mg/ ml, and stored at -80 °C after flash-frozen with liquid nitrogen.

### TEV protease expression and purification

The tobacco etch virus (TEV) encoded by the plasmid pRK792^64^ was overexpressed in a tRNA-supplemented BL21-CodonPlus (DE3)-RIPL strain. The cells were grown in LB broth containing 100 µg/mL ampicillin and 30 µg/mL chloramphenicol at 30 °C until *A*_600_ of 0.8 ∼1.0. At this point, the temperature was decreased to 18 °C, and protein expression was induced by 0.5 mM IPTG overnight. The cells in 50 mM KPi, pH 8.0, 500 mM NaCl, and 10% glycerol were lysed by a microfluidizer and the supernatants after removing the cell debris were subjected to affinity chromatography using Talon resin. The eluted proteins were concentrated to ∼3 mg/mL and quickly dialyzed against 20 mM Tris-HCl, pH 7.5, 200 mM NaCl, and 10% glycerol for 3 - 4 hours, and stored at -80 °C after flash-frozen with liquid nitrogen. A typical yield can reach >30 mg/L.

### MelB_St_ reconstitution into lipids nanodiscs

The MelB_St_ reconstitution was performed as described previously^28^. Briefly, 5.6 mg of *E. coli* polar lipids extract at a concentration of 40 mg/mL solubilized in 7.5% DDM was added to 1 mL of 1 mg of MelB_St_ in 20 mM Tris-HCl, pH 7.5, 100 mM NaCl, 10% glycerol, and 0.35% UDM (∼1:350 mole/mole), and incubated for 10 min on ice. MSP1E3D1 proteins were then added into the MelB_St_/lipid mixture at a 5:1 mole ratio (MSP1E3D1:MelB_St_), and incubated at room temperature with mild stirring for 30 min before being shifted to 4 °C with mild stirring. The detergents were removed and subsequent MelB_St_ lipids nanodiscs were formed by stepwise additions of Bio-beads SM-2 resin (500 mg and then an additional 300 mg after 2 h) and further incubated overnight. After separating the Bio-beads SM-2 resins, the MelB_St_ lipids nanodiscs were isolated by absorbing onto Ni-NTA beads and the MelB_St_-free empty nanodiscs were in the void due to His-tag being removed. The MelB_St_ lipids nanodiscs were eluted by adding 250 mM imidazole and the elutes were dialyzed against 20 mM Tris-HCl, pH 7.5, and 150 mM NaCl, and concentrated to ∼ 3-4 mg/mL. After ultracentrifugation at ∼352000 g (90,000 rpm using a TLA-120 rotor) in a Beckman OptimaTM MAX Ultracentrifuge for 30 min 4 °C, the supernatant containing MelB_St_ lipids nanodiscs were stored at -80 °C after flash-frozen with liquid nitrogen.

### Nbs expression and purification

*E. coli* BL21(DE3) Arctic express cells were used to produce Nbs. The Nb725_4 and anti-Fab Nb were expressed by the expression plasmid pET26/Nb725_4 and pET26/anti-Fab Nb, respectively, containing the *pelB* leader sequence. Nb725 was expressed by the plasmid p7xC3H/Nb725 as described^29^. Cells were grown in Terrific Broth media supplemented with 0.4% (vol/vol) glycerol at 37 °C. Protein expression was induced by adding 1 mM IPTG at the mid-log phase after decreasing the temperature to 25 °C, and the cultures continued growth for 18 to 20 h. The periplasmic extraction of Nbs using sucrose-mediated osmolysis as described^65^, and Nb purification was performed by cobalt-affinity chromatography, which yielded ∼5 mg/L with >95% purity. The cytoplasmic expression for the Nb725 also provided a similar yield and purity, and all were stable. The purified Nbs were concentrated to ∼5-10 mg/mL, dialyzed against 20 mM Tris-HCl, pH 7.5, and 150 mM NaCl, and frozen in liquid nitrogen and stored at -80 °C.

### NabFab expression and purification

The NabFab expressing plasmid pRH2.2/NabFab and the expression protocol were kindly gifted from Dr. Kasper P. Locher^36,66^. Briefly, the NabFab was expressed in *E. coli* C43 (DE3) cells in an autoinduction medium at 25 °C for ∼18 h. The cells were resuspended in 20 mM Tris-HCl, pH 7.5, 500 mM NaCl, and 1 mM EDTA, and lysed by a microfluidizer. The lysates were subjected to heat treatment at 65 °C for 30 min, followed by centrifugation at 33175 g (15,000 rpm on the rotor JA18) for 30 min at 4 °C to remove insoluble materials. The supernatants were loaded onto the 5-mL Cytivia Protein L column for affinity purification of NabFab proteins. The eluted NabFab in 0.1 M acetic acid, pH 2.0, was neutralized with 200 mM Tris-HCl, pH 8.5, dialyzed against 20 mM Tris-HCl, pH 7.5, 150 mM NaCl, and concentrated up to 5 mg/mg prior to being frozen in liquid nitrogen and stored at -80 °C. The yield for NabFab protein is about 2-3 mg/L.

### EIIA^Glc^ expression and purification

The overexpression of unphosphorylated EIIA^Glc^ was performed in the *E. coli* T7 Express strain (New England Biolabs) as described^33^.

### A bacterial two-hybrid assay

A bacterial two-hybrid assay based on reconstituting adenylate cyclase activities from the T18/T25 fragments of *Bordetella pertussis* adenylate cyclase toxin as described^67,68^ was used to examine *in vivo* interaction of Nb and MelB_St_ in *E. coli* DH5α *cyaA* strain^29^. The competent cells were transformed by two compatible plasmids with different replication origins (pACYC/T25:MelB_St_ and pCS19/Nb725:T18 or pCS19/Nb725-_4:T18)^69^ containing two hybrids (pACYC)^70^ and plated onto the lactose-free MacConkey agar plate containing 30 mM maltose as the sole carbohydrate source, 100 mg/L ampicillin, 25 mg/L chloramphenicol, and 0.2 mM IPTG, and the plates were incubated in 30 ℃ for 7 days before photography. Red colonies grown on the MacConkey agar indicated positive fermentation due to the cAMP production derived from the interaction of two hybrids. Yellow colonies denoted no fermentation and no cAMP production.

### Sugar fermentation assay

*E. coli* DW2 cells (Δ*melBΔlacYZ)* were transformed with two compatible plasmids with different replication origins (pACYC/MelB_St_ and pCS19/Nbs)^70^ and plated onto the lactose-free MacConkey agar plate containing 30 mM maltose or melibiose as the sole carbohydrate source, 100 mg/L ampicillin, 25 mg/L chloramphenicol, and 0.2 mM IPTG. The plates were incubated at 37 ℃ for 18 h before photography. Magenta colonies, a normal sugar fermentation; yellow colonies, no sugar fermentation. The cells carrying two empty plasmids or with MelB_St_ but no Nb were used as the control. Maltose fermentation was another control for the specificity of Nb inhibition.

### [^3^H]Melibiose transport assay and western blot

*E. coli* DW2 cells (*mel*A^+^, *melB^-^, lacZ^-^Y^-^*)^71^ were transformed with the two compatible plasmids pACYC/MelB_St_ and pCS19/Nb725 or pCS19/Nb725_4, respectively. The cells transformed with two empty vectors (pACYC and pCS19) or pACYC/MelB_St_ with pCS19 without an Nb were used as the negative or positive control, respectively. The cells were grown in LB broth with 100 mg/L ampicillin and 25 mg/L chloramphenicol in a 37 °C shaker overnight, and inoculated by 5% to fresh LB broth with 0.5% glycerol, 100 mg/L ampicillin, 25 mg/L chloramphenicol, and 0.2 mM IPTG. The cultures were shaken at 30 °C for 5 h. The transport assay was performed at 20 mM Na^+^ and 0.4 mM [^3^H]melibiose (specific activity of 10 mCi/mmol) in 100 mM KP_i_, pH 7.5, 10 mM MgSO_4_ at *A*_420_ of 10 (∼0.7 mg proteins/mL) as described^25,56^. The transport time courses were carried out at zero, 5 s, 10 s, 30 s, 1 m, 2 m, 5 m, 10 m, and 30 m by dilution and fast filtration. The filters with trapped cells were subjected to radioactivity measurements using a Liquid Scintillation Counter. The MelB_St_ membrane expression was analyzed by Western blot using anti-His tag antibody as described^29^.

### Isothermal titration calorimetry

All ITC ligand-binding assays were performed with the Nano-ITC device (TA Instruments)^17^ and the heat releases are plotted as positive peaks. All samples were buffer-matched by simultaneously dialyzed in the same buffer, or diluting the concentrated small molecule ligands to corresponding dialysis buffers as described^17,33^. The titrand (MelB_St_) placed in the ITC Sample Cell was titrated with the specified titrant (placed in the Syringe) by an incremental injection of 1.5 **-** 2 μL aliquots at an interval of 300 sec at a constant stirring rate of 250 rpm. All samples were degassed by a TA Instruments Degassing Station (model 6326) for 15 min prior to the titration. Heat changes were collected at 25 °C, and data processing was performed with the NanoAnalyze (v. 3.7.5 software) provided with the instrument. The normalized heat changes were subtracted from the heat of dilution elicited by the last few injections, where no further binding occurred, and the corrected heat changes were plotted against the mole ratio of titrant versus titrand. The values for the binding association constant (*K*_a_) were obtained by fitting the data using the one-site independent-binding model included in the NanoAnalyze software (version 3.7.5), and the dissociation constant (*K*_d_) = 1/*K*_a_. In most cases, the binding stoichiometry (*N*) number was fixed to 1.

### Protein concentration assay

The Micro BCA Protein Assay (Pierce Biotechnology, Inc.) was used to determine the protein concentration assay. The *A*_280_ was used for concentration estimation.

### Protein complex preparation for grid vitrification

Proteins NabFab, Nb725_4, anti-Fab Nb were mixed at a mole ratio of 1:1.1:1.15. The final complex was obtained by adding WT MelB_St_ in nanodiscs based on a mole ratio of 1:1.6 of MelB_St_:NabFab complex. The sample was expected to contain 6 μM MelB_St_, 12 μM MPS1E3D1, 9.5 μM NabFab complex in 20 mM Tris-HCl, pH 7.5, 150 mM NaCl (∼1.5 mg/mL). The complex was also analyzed by gel-filtration chromatography using GE Superdex 200 column on a BioRad NGC Chromatography System. Peak fractions were examined by SDS-15% PAGE stained by silver nitrate.

### Grids vitrification and single-particle cryoEM data acquisition

Grids preparation and data collection were carried out in S^2^C^2^ (Menlo Park, CA). The WT MelB_St_/Nb725_4/NabFab/anti-NabFab Nb complex was thawed out from -80 °C storage. Aliquots of 3 μL of samples at 0.75 mg/ml or 1.5 mg/ml protein concentration were placed on 300-mesh Cu holey carbon grids (Quantifoil, R1.2/1.3) after glow-discharge (PELCO easiGlow™) at 15 mA for 30 sec, and blotted on two sides for 3 sec at 4 °C and 100% humidity before vitrified in liquid ethane using a Vitrobot Mark IV (Thermo Fisher). The grids were subsequently transferred to a Titan Krios (G3i) electron microscope (Thermo Fisher) operating at 300 kV and equipped with K3 direct electron detector (Gatan) and BioQuantum energy filter. The best grids were found from the sample at 1.5 mg/ml concentration.

A total of 14,094 and 8715 movies were automatically collected using the EPU Data Acquisition Software (Thermo Fisher) for non-tilted and 30-degree tilted collections, respectively. Both datasets were collected at a 0.86 Å pixel size and a dose of 50 e^−^/Å^−2^ across 40 frames (0.0535 or 0.07025 sec per frame for the non-tilted and tilted data collection, respectively) at a dose rate of approximately 23.36 or 17.8 e^−^/Å^2^/s, respectively. A set of defocus values was applied ranging from -0.8, -1.0, -1.2, -1.4, or -1.8 μm with an energy filter slit width of 20 eV and a 100 μm objective aperture.

### Single-particle data processing

All cryoEM single-particle data processing was performed using CryoSPARC program (v. 2 – 4.01) (**sFig. 4a**). All imported movies were aligned using defaulted Patch Motion Correction and followed with defaulted Patch contrast transfer function (CTF) Estimation. Micrographs were then curated based on CFT Estimation, astigmatism, and ice thickness, which resulted in 13,649 and 7,129 micrographs for the non-tilted and 30-degree tilted datasets, respectively.

Initially, particle picking from the 2000 micrographs was performed using Blob Picker using 120 and 160 Å for the minimal and maximal particle diameters, respectively, in the CryoSPARC program (**sFig. 4b)**. A total of 1,087,124 particles were selected and extracted with a box size of 256 pixels. After 2 runs of 2D Classification, five classes were selected as imputing templates in Template Picker. A total of resultant 9,103,229 particles from 13,649 micrographs were extracted with a box size of 384 pixels bin 92 pixels and subjected to iterative rounds of 2D Classification. The yielded 1,245,734 particles were re-extracted at a box size of 416 pixels. After further sorting using 2D Classification, the highly selected 338,883 particles, which clearly contained nanodisc-surrounding MelB_St_ bound with Nb725_4/NabFab/anti-NabFab Nb, were used for Ab-Initio 3D Reconstruction. One out of 5 Ab-Initio classes contained the expected super-complex from a total of 93,693 particles, which allowed for a 3.80 Å volume to be reconstructed using Non-uniform Refinement. The 93,693 particles were then further sorted using heterogeneous refinement with input volumes created by multiclass Ab-initio to a subset of 41,264 particles, which allowed a 3.49 Å volume to be reconstructed using Non-uniform Refinement with a static mask prepared from one of the volumes. This map was successfully used to build the initial model of all four proteins; however, this map was anisotropic and lacked some orientations in angular space. To improve the map quality, two strategies were applied including reprocessing this dataset and performing tilted data collection. The hand of the 3.49 Å volume was changed and used to generate 50 templates.

Template Pick using the 50 templated was applied to the 13,649 micrographs, which generated 7,632,727 particles (**sFig. 4c)**. After extraction at a box size of 384 pixels bin 92 pixels and iterative rounds of 2D Classification and Heterogeneous Refinement, 370,941 particles were selected and re-extract at a box size of 384 pixels. This 370,941-particle set was combined with the previously identified 41,264-particle set after extraction at a box size of 384 pixels and removed duplicate, the subset of 188,566 particles exhibited slightly improved directional distribution. To further explore the first data collection, the previously rejected 2D classes were analyzed and 214,522 particles were selected based on orientation. After combined and iterative rounds of 2D Classification, Heterogeneous Refinement, and Local Refinement, a subset of the 196,254-particle set supported a 3.39 Å volume to be reconstructed using local refinement, and the directional distribution of the map is further improved.

A tilted data collection on the same microscopy and parameter settings was performed with a 30-degree tilt (**sFig. 4c)**. A total number of 7,129 micrographs were obtained. The 50-templates were used for particle pick, which yielded 2,887,147 particles after curation. The particles were extracted using a box size of 384 bin 96 pixels. After 3-run 2D Classification and followed 3-run Heterogeneous Refinement using the 3.39 Å volumes and the other two Ab-initio volumes, a subset of 544,531 particles were re-extracted using a box size of 384 pixels. After further iterative rounds of 2D Classification, Heterogeneous Refinement, and Local Refinement, 182,101 particles were obtained for the tilted collection, which could support a 4.01 Å volume using Local Refinement and directional distribution focused on the missing angles in the first dataset.

A total number of 378,355 polished particles from both non-tilted and 30-degree tilted data acquisition were used to reconstruct a 3.40 Å volume using Local Refinement, and the 30-degree tilted data filled the missing particle orientations but the resolution was not improved. The particles were subjected to 3D Classifications using a target resolution of 3.3 Å and 10 classes and the highest resolution classes were collected. The final collected 296,925-particle set supported a 3.37 Å volume to be reconstructed using Local Refinement and a static mask covering MelB_St_, Nb725_4, and NabFab. The reporting GSFSC resolution of 3.29 Å was calculated by Validation (FSC) using the two half maps, and the particle distribution was calculated by ThereDFSC (**sFig. 5**).

The initial mask for Non-uniformed Refinement was created in UCSF ChimeraX using the volume generated from Ab-Initio 3D Reconstruction which covered the full complex. The refinement masks for final-stage Local Refinement only covered MelB_St_, Nb725_4, and NabFab and filtered at a low pass to 10 Å and soft padding width of 10. The 3D Classification used the full-covered mask as the solvent mask and the refinement masks in the Local Refinement as the focus mask.

Local Refinement set the initial low pass resolution of 8 Å, use of pose/shift Gaussian prior during alignment, Rotation search extent of 9 °, and Shift search extent of 6 Å. The final 3.29 Å volume was sharpened in Phenex auto_sharpen program^72^ default setting for model refinement.

### Model building and structure refinement

The initial model building was carried out based on the 3.45 Å reconstruction map obtained from the first dataset. The coordinates of MelB_St_ D59C (PDB ID 7L17), NabFab and anti-NabFab Nb (PDB ID 7PHP), and a sequence-based 3-D model generated by AlphaFold 2 were fitted into the map in UCSF ChimeraX^73^. The N-terminal helices of MelB_St_ fit well and the C-terminal helices were manually fitted into the initial map. The sequence of MelB_St_ D59C was mutated back to wild type in COOT^74^. All 4 chains were changed to poly-Ala peptides and the side chains of each residue were rebuilt. Structure refinement was performed using phenix.real_space_refine application in PHENIX^75^ in real space with secondary structure and geometry restraints.

The model was refined against the final map at a resolution of 3.29 Å after sharpening using PHENIX auto_sharpen program default setting^72^. A cation Na^+^ was modeled in the density between helices-II and -IV in COOT. After runs of refinement and remodeling, in total, 417 residues of MelB_St_ (positions 2-210, 219-355, and 364-432), with 6 unassigned side-chains at the C-terminal domain (Leu293, Tyr355, Arg363, Tyr369, Tyr396, and Met410) due to the map disorder, 122 residues of Nb725_4 (2-123), 229 residues of NabFab H-chain (1-214), and 210 residues of NabFab L-chain (4-213), were modeled, respectively. There are three MelB_St_ regions with missing densities including the middle loop righter after the Nb binding site (positions 211-218), loop_10-11_ (356-361), and the C-terminal tail after residue Tyr432. In addition, the densities between positions 219-230 and the regions close to the missing densities at loop_10-11_ are very weak. There is no unexplained density that can be clearly modeled by lipids. Statistics of the map reconstruction and model refinement can be found in **sTable 1**. The model-map fitting quality was evaluated by *Q* score program^42^. Pymol^76^ and UCSF chimera or ChimeraX^73^ were used to prepare the figures.

### All-atom molecular dynamics (MD) simulations

All-atom MD simulations were performed taking the inward-facing CryoEM structure as the starting model. Atom coordinates in the loops missing from the CryoEM structure were filled in using homology modeling with the Modeller software package^77^. Subsequently, the protein was embedded into a lipid bilayer composed of 1-palmitoyl-2-oleoyl phosphatidylethanolamine (POPE), 1-palmitoyl-2-oleoyl phosphatidylglycerol (POPG), and cardiolipin (CDL), in a molar ratio of 7:2:1 for POPE:POPG:CDL, mimicking the membrane composition of *E. coli.* Each side of the lipid bilayer was then enclosed by a box of water molecules with 25 Å thickness. Sodium chloride ions were introduced among these water molecules to yield a solution with an approximate concentration of 0.16 M. The system setup was performed via the CHARMMGUI^78^ web interface. The all-atom MD simulations for the outward-facing MelB_St_ under the same setup have been reported by constructing through an in-silico mutation, converting the outward-facing crystal structure of the D59C MelB_St_ mutant (PDB ID 7L16) back to the WT MelB_St_^44^. In both the inward and outward-facing configurations, a sodium ion was initially situated in the binding site, while melibiose was notably absent from the system. Each system underwent initial optimization where the heavy atoms of proteins and lipids were harmonically restrained, employing a force constant of 1000 kJ/mol/Å^2^. This was followed by 100 picoseconds (ps) of dynamics at a temperature of 300 K in the constant NVT ensemble. Subsequently, the restraints were gradually reduced during 10 ns of dynamics in the constant NPT ensemble, maintained at a temperature of 300 K and pressure of 1 atm. For equilibration, we carried out an additional 30 ns of dynamics in the NPT ensemble under the same conditions, but without any restraints. After the equilibration stage, 1 μs of dynamics was performed in the constant NPT ensemble, under the same temperature and pressure, to generate the production trajectories. We maintained the temperature using a Langevin thermostat with a friction coefficient of 1 ps^-1^. The trajectory was propagated with a timestep of 2 femtoseconds (fs), and all bonds involving a hydrogen atom were constrained. The CHARMM36 force field^79–81^ was utilized to simulate the protein, lipid, and ions throughout all simulations. Electrostatic interactions were calculated using the particle mesh Ewald method, with a tolerance of 5 x 10^-4^. We set the cutoff for van der Waals interactions at 12 Å. The TIP3P model^82^ was used to treat water molecules and ions. All classical MD simulations were performed using the OPENMM^83^ software package.

### Hydrogen-deuterium exchange coupled to mass spectrometry (HDX-MS)

In-solution pairwise HDX-MS experiment was performed to study the structural dynamics of MelB_St_. Labeling, quenching, lipids removal, and online digestion were achieved using a fully automated manner using HDx3 extended parallel system (LEAP Technologies, Morrisville, NC)^84,85^.

Working samples of MelB_St_ and MelB_St_ complexed with Nb725_4 were prepared to a concentration of 50.0 μM for MelB_St_ and 100 μM for Nb725_4 in a buffer of 25 mM Tris-HCl, pH 7.5, 150 mM NaCl, 10% glycerol, 0.01% DDM in H_2_O. Aliquots of 4 µl of each working sample were diluted by 10-fold into the labeling buffer of 25 mM Tris-HCl, pD 7.5, 150 mM NaCl, 10% glycerol, 0.01% DDM in D_2_O. The reactions were incubated in D_2_O buffer at 20 °C for multiple time points (30 sec, 300 sec, and 3000 sec) in triplicates and non-deuterated controls were prepared in a similar manner except H_2_O buffer was used in the labeling step. The pH of the labeling buffer was measured with a pH meter mounted with a glass electrode and then corrected to pD (pD = pH + 0.4).

The reactions were quenched at given time points by adding the same volume of ice-cold 6 M urea, 100 mM citric acid, pH, 2.3 in water for 180 seconds at 0 °C and immediately subjected to a lipid filtration module containing ZrO_2_ available on the LEAP PAL system. After incubation of 60 sec, the LEAP X-Press then compressed the filter assembly to separate proteins from the ZrO2 particles-bound phospholipids and detergents. The filtered protein sample was injected into a cool box for online digestion and separation.

LC/MS bottom-up HDX was performed using a Thermo Scientific™ Ultimate™ 3000 UHPLC system and Thermo Scientific^TM^ Orbitrap Eclipse^TM^ Tribrid^TM^ mass spectrometer. Samples were digested with a Nepenthesin-2 (Affipro, Czech Republic) column at 8 °C for 180 sec and then trapped in a 1.0 mm x 5.0 mm, 5.0 µm trap cartridge for desalting. Peptides were then separated on a Thermo Scientific™ Hypersil Gold^TM^, 50 x1 mm, 1.9 µm, C18 column with a gradient of 10 % to 40 % gradient (A: water, 0.1 % formic acid; B: acetonitrile, 0.1 % formic acid) gradient for 15 minutes at a flow rate of 40 µL/min. To limit carry-over issues, a pepsin wash was added in between runs. To limit the back-exchange of hydrogen, all of the quenching, trapping, and separation steps were performed at near 0 °C.

For data analysis, a nonspecific digested peptide database was created for MelB_St_ with separate tandem mass spectrometry measurements of non-deuterated samples. Digested peptides from undeuterated MelB_St_ protein were identified on the orbitrap mass spectrometer using the same LC gradient as the HDX-MS experiment with a combination of data-dependent and targeted HCD-MS2 acquisition. Using the Thermo BioPharma Finder software (v 5.1), MS2 spectra were matched to the MelB_St_ sequence with fixed modifications. Altogether 153 peptide assignments (confident HDX data cross all labeling times) were confirmed for MelB _St_ samples giving 86% sequence coverage. MS data files were processed using the Sierra Analytics HDExaminer software with the MelB_St_ peptide database. Following the automated HDX-MS analysis, manual curation was performed. Upon the completion of the data review, a single charge state with high-quality spectra for all replicates across all HDX labeling times was chosen to represent HDX for each peptide. Differential HDX data were tested for statistical significance using the hybrid significance testing criteria method with an in-house MATLAB script, where the HDX differences at different protein states were calculated (ΔD = D_Nb725_4-bound_ – D_Nb-free_). D denotes deuteration, defined as the mass increase above the average mass of the undeuterated peptide. Mean HDX differences from the three replicates were assigned as significant according to the hybrid criteria based on the pooled standard deviation and Welch’s t-test with P < 0.01. The statistically significant differences observed at each residue (ie., ΔD ≥ 0.3184 for this study) were used to map HDX consensus effects based on overlapping peptides onto the structure models. Residue-level data analysis was performed using a built-in function in BioPharma Finder software. The parameters included the number of simulations of 200, recorded solutions of 20, Chi2 increase by the larger of smooth absolute of 0, smother relative of 2, differential absolute of 0, and differential relative of 2.

## Competing Interests

Y. S. and R. V. are employees of *Thermo Fisher Scientific*.

## Acknowledgments

We thank the S^2^C^2^ cryoEM facility sponsored by NIH Common Fund Transformative High-Resolution Cryo-Electron Microscopy Program (U24 GM129541). We thank Drs. Wah Chiu, Htet Khant, and Grigore Pintilie for support and critical discussions; Drs. Kaspar P. Locher, Anthony A. Kossiakoff, Joël S. Bloch, and Somnath Mukherjee for kindly providing the NabFab plasmid and detailed instructions; Drs Eric R. Geertsma and Raimund Dutzler for the FX cloning tools; Dr. Gerard Leblanc for a MelB-expressing vector and DW2 strain; Drs. Bryan Sutton and Kei Nanatani for support; and Dr. Michael Wiener for critical reading and editing. This work was supported by the National Institutes of Health Grant R01 GM122759 to L.G. and the Welch Foundation D-2108-20220331 to R. L.

## Notes

### Summary of Updates

While many 3D structures of cation-coupled transporters have been determined, the mechanistic details governing the obligatory coupling and functional regulations still remain elusive. The bacterial melibiose transporter (MelB) is a prototype of the Na+-coupled major facilitator superfamily transporters. With a conformational nanobody (Nb), we determined a low-sugar affinity inward-facing Na+-bound cryoEM structure. Collectively with the available outward-facing sugar-bound structures, both the outer and inner barriers were localized. The N-and C-terminal residues of the inner barrier contribute to the sugar selectivity pocket. When the inner barrier is broken as shown in the inward-open conformation, the sugar selectivity pocket is also broken. The binding assays by isothermal titration calorimetry revealed that this inward-facing conformation trapped by the conformation-selective Nb exhibited a greatly decreased sugar-binding affinity, suggesting the mechanisms for the substrate intracellular release and accumulation. While the inner/outer barrier shift directly regulates the sugar-binding affinity, it has little or no effect on the cation binding, which is also supported by molecular dynamics simulations. Furthermore, the use of this Nb in combination with the hydron/deuterium exchange mass spectrometry allowed us to identify dynamic regions; some regions are involved in the functionally important inner barrier-specific salt-bridge network, which indicates their critical roles in the barrier switching mechanisms for transport. These complementary results provided structural and dynamic insights into the mobile barrier mechanism for cation-coupled symport.

