## Supplemental materials for "Mobile barrier mechanisms for Na^+^-coupled symport in an MFS sugar transporter"

**Contents:**

- **Supplemental Table**
  sTable 1 Cryo-EM data collection and structure determination statistics.
  sTable 2 HDX reaction and labeling details.
  sTable 3. *E. coli* strains and plasmids used in this study.

- **Supplemental Figures**
  sFig. 1. Hybrid Nb725_4 generated by CDR grafting.
  sFig. 2. Effects of substrate/ligand binding on the Nbs binding.
  sFig. 3. Nb effects on substrate/ligand binding.
  sFig. 4. CryoEM data process.
  sFig. 5. GSFSC resolution and 3dFSC.
  sFig. 6. Evaluation of map and models.
  sFig. 7. NabFab comparison.
  sFig. 8. Interactions of Nb725m_4 and $MelB_{St}$.
  sFig. 9. Complex of $MelB_{St}$ with Nb725 and $EIIA^{Glc}$.
  sFig. 10. MD simulations of the $Na^+$ binding at both inward- and outward-facing states.
  sFig. 11. Galactose-binding pocket in the outward-facing crystal structure [PDB ID 7L17].
  sFig. 12. Alignment.
  sFig. 13. Membrane topology.
  sFig. 14. Histograms of deuterium uptake time courses.

**sTable 1. Cryo-EM data collection and structure determination statistics.**

| | $MelB_{St}$/Nb725m/NabFab | |
|---|---|---|
| **EMDB Accession Code** | EMD-41062 | |
| **PDB Accession ID** | 8T60 | |
| | Non-tilted collection | Tilted collection |
| **Data Collection** | | |
| Microscope | Krios-TEMBETA | Krios- TEMBETA |
| Voltage (kV) | 300 | 300 |
| Number of movies | 14,094 | 8,716 |
| Electron dose ($e^-/Å^2$) | 50.00 | 50.00 |
| Defocus range (μm) | −0.8 to -1.8 | −0.8 to -1.8 |
| Pixel size (Å) | 0.86 | 0.86 |
| Plate tilt angel (Degree) | / | 30 |
| **Data Processing** | | |
| Initial number of particles | 7,632,727 | 2,887,147 |
| Combined final number of particles | 203,876 | |
| Symmetry imposed | C1 | |
| Map resolution* (Å) | 3.29 | |
| B factor | 101.4 | |
| **Model Refinement** | | |
| Chains | 5 | |
| Non-hydrogen atoms | 7503 | |
| Protein residues | 978 | |
| Mean B-factor | | |
| Protein | 97.35 | |
| $Na^+$ ion | 88.64 | |
| RMS deviations | | |
| Bond lengths (Å) | 0.003 | |
| Bond angles (°) | 0.509 | |
| MolProbity score | 1.82 | |
| Clash score | 4.09 | |
| Poor rotamers (%) | 1.98 | |
| Ramachandran plot | | |
| Favored (%) | 93.48 | |
| Allowed (%) | 6.52 | |
| Outliers (%) | 0.00 | |
| Model Resolution (Å)† | 3.2/3.3/3.5 | |

*Resolution determined by Fourier shell coefficient threshold of 0.143 for corrected masked map
† Resolution determined between the model and the resolved map by Fourier shell coefficient threshold of 0/0.143/0.5

#### sTable 2. HDX reaction and labeling details.

| Samples measured | WT $MelB_{St}$ | WT $MelB_{St}$ complexed with Nb725m |
|---|---|---|
| HX reaction buffer | 25 mM Tris-HCl, pD 7.5, 150 mM NaCl, 10% Glycerol, and 0.01% DDM | |
| Reaction temperature (°C) | 20 | |
| HX time course (s) | 0, 30, 300, 3000 | |
| Number of peptides | 153 | |
| Sequence coverage by labeling | 86 % | |
| Mean peptide length | 9.3 | |
| Average redundancy | 2.9 | |
| Replicates (technical) | 3 | |
| $\|\Delta D\|$ (Da) | 0.3071 | |
| Back exchange rate | Not appliable | |

#### sTable 3. *E. coli* strains and plasmids used in this study

| | *Description* | Reference |
|---|---|---|
| **Stains** | | |
| *E. coli* DW2 | *melA*$^{+}$ Δ*melB* Δ*lacZY* | 59 |
| *E. coli* XL1 Blue | *recA1 endA1 gyrA96 thi-1 hsdR17 supE44 relA1 lac [F' proAB lacIqZΔM15 Tn10 (Tetr)]* | Agilent Technologies |
| *E. coli* ArcticExpress (DE3) | $F^{-}$ *ompT hsdS*($rB^{-}$ $mB^{-}$) *dcm*$^{+}$ Tetr gal λ(DE3) *endA* Hte [cpn10 cpn60 Gentr ] | Agilent Technologies |
| *E. coli* DH5α *cyaA*$^{-}$ | Δ*cyaA* | 29 |
| *E. coli* BL21(DE3) T7 express | *fhuA2 lacZ::T7 gene1 [lon] ompT gal sulA11 R(mcr-73::miniTn10*--Tet$^{S}$*)2 [dcm] R(zgb-210::Tn10*--Tet$^{S}$*) endA1 Δ(mcrC-mrr)114::IS10* | New England Biolabs. |
| *E. coli* BL21(DE3) C43 | $F^{-}$ *ompT hsdSB (rB- mB-) gal dcm (DE3)* | 60 |
| *E. coli* BL21(DE3) pRIL | $F^{-}$ ompT hsdS($rB^{-}$ $mB^{-}$) dcm$^{+}$ Tet$^{r}$ gal endA Hte [argU ileY leuW], Cam$^{r}$ | Agilent Technologies |
| **Plasmids** | | |
| pCS19 | pQE60 derivative inserted with gene *lacI*$^{q}$; amp$^{r}$ | 69 |
| pACYC | pACYC/FX-derived vector for control; no *ccdB* gene: cam$^{r}$ | 70 |
| pCS19/FX | Expression vector with two SapI sites and *ccdB* gene for FX cloning; amp$^{r}$ | 70 |
| pACYC/$MelB_{St}$ | Expression vector for $MelB_{St}$ derived from pACYC/FX; cam$^{r}$ | 70 |
| pCS19/X:T18/FX | Expression plasmid for expressing a target protein "X" with a C-terminal fusion with T18 fragment; two SapI sites and *ccdB* gene for FX cloning; amp$^{r}$ | 29 |
| pCS19/T18 | pCS19/X:T18/FX-derived vector for expressing T18 fragment; no *ccdB* gene. | 29 |
| pACYC/T25 | pACYC/T25:X/FX-derived vector for expressing T25 fragment; no *ccdB* gene | 29 |
| pACYC/T25:$MelB_{St}$ | Expression plasmid for the T25:$MelB_{St}$ hybrid derived from pACYC/T25:X/FX; cam$^{r}$ | 29 |
| pCS19/Nb725:T18 | Expression plasmid for Nb725:T18 hybrid derived from pCS19/X:T18/FX; amp$^{r}$ | 29 |
| pCS19/Nb725_4:T18 | Expression plasmid for Nb728:T18 hybrid derived from pCS19/X:T18/FX; amp$^{r}$ | This study |
| pCS19/Nb725 | Expression plasmid for Nb725 derived from pCS19/FX; amp$^{r}$ | 29 |
| pCS19/Nb725_4 | Expression plasmid for Nb725_4 derived from pCS19/FX; amp$^{r}$ | This study |
| pET26b(+) | Construction of Nb725_4 expression plasmid, construction of anti-Fab Nb expression plasmid; replacement pelB leader sequence for periplasmic expression, Kan$^{r}$ | Novagen (EMD Millipore) |
| pET26/Nb725_4 | Inducible periplasmic expression plasmid used for Nb725_4 protein production, Kan$^{r}$ | This study |
| pET26/Anti-Fab Nb | Inducible periplasmic expression plasmid used for Anti-Fab Nb protein production, Kan$^{r}$ | This study |
| p7XC3H/Nb725 | Inducible cytoplasmic expression plasmid used for Nb725 protein production, Kan$^{r}$ | 29 |

| | | |
|---|---|---|
| p7XNH3/EIIA$^{Glc}$ | Inducible cytoplasmic expression plasmid used for *E. coli* EIIA$^{Glc}$ protein production, Kan$^{r}$ | 33 |
| pR2.2/NabFab | Expression and purification of NabFab, Amp$^{r}$ | 36 |
| pMSP1E3D1 | Expression and purification of MSP1E3D1, Kan$^{r}$ | Addgene/20066 |
| pRK792 | Expression and purification of TEV protease, Amp$^{r}$ | Addgene/8830 |

#### Primers

| Labels | Applications | Oligonucleotides |
|---|---|---|
| MelB_Nb | pCS19/Nb725_4:T18 (This adds C-terminal 'A' and allow the C-terminal fusion) | Fwd: 5′- ATATATGCTCTTCTAGTCAACGTCAATTGGTAG -3′ |
| | | Rev: 5′- TATATAGCTCTTCATGCGCTGCTCACGGTCAC -3′ |
| MelB_FX | pCS19:Nb725_4-CTH (This adds C-terminal 'HHHHHH and truncated by a stop codon) | Fwd: 5′- ATATATGCTCTTCTAGTATGCAACGTCAATTGGTAG -3′ |
| | | Rev: 5′- TATATAGCTCTTCATGCTTAGTGGTGATGATGGTGGTGGCT GCTCACGGTCAC -3′ |
| Nb-PelB-NdeI | Add restriction site for Nb725_4 and Anti-Fab Nb construction | Fwd: 5'-TTTAAGAAGGAGATATACATATG-3' |
| Nb-STRP-XhoI | Add restriction site for Nb725_4 and Anti-Fab Nb construction | Rev: 5'TTTGTTCTAGACTCGAGTTATTTCTC-5' |

#### Supplementary figures

**sFig. 1**

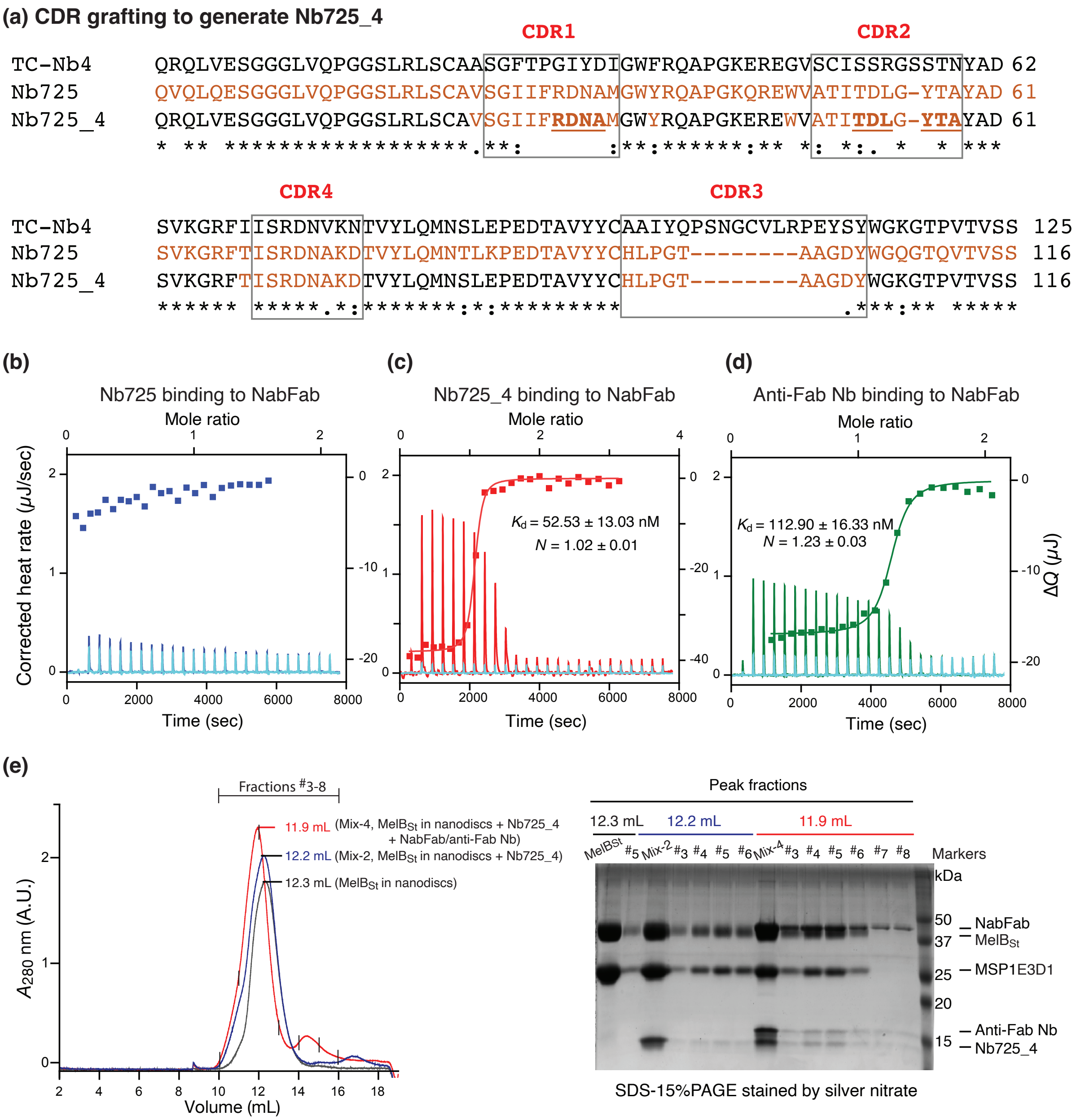

**sFig. 1. Hybrid Nb725_4 generated by CDR grafting. (a)** All CDR regions are indicated by boxes. The sequences of TC-Nb4 and $MelB_{St}$ Nb725 were colored in black and red, respectively. The sequence of hybrid Nb725_4 was colored in black and red from the two parents Nbs. The Nb725_4 binding residues were highlighted in underlined bold. **(b-d)** ITC measurements of Nbs binding to the NabFab. The $K_d$ values for Nb725_4 and the Anti-Fab Nb binding to NabFab were presented as the mean ± sem and the number of tests = 2. $N$, the binding stoichiometry number. The interaction of the parent Nb725 to the NabFab was too weak to be fit accurately. **(e)** Isolation of the $MelB_{St}$/Nb725_4/NabFab/anti-Fab Nb complex. $MelB_{St}$ proteins were reconstituted into the

lipid nanodiscs with the membrane scaffold protein 1E3D1 (MSP1E3D1). Sample containing $MelB_{St}$ (black curve), $MelB_{St}$ with Nb725_4 (blue curve), or with Nb725_4, NabFab, and anti-Fab Nb (red curve) were prepared prior to analysis by gel filtration chromatography in a combination of the SDS-15%PAGE. The mix-2 sample, $MelB_{St}$ with Nb725_4, blue color; mix-4 sample, $MelB_{St}$ with Nb725_4, NabFab and anti-Fab Nb, red color. The mixture of each protein in the loading buffer does not contain a reducing agent.

sFig. 2

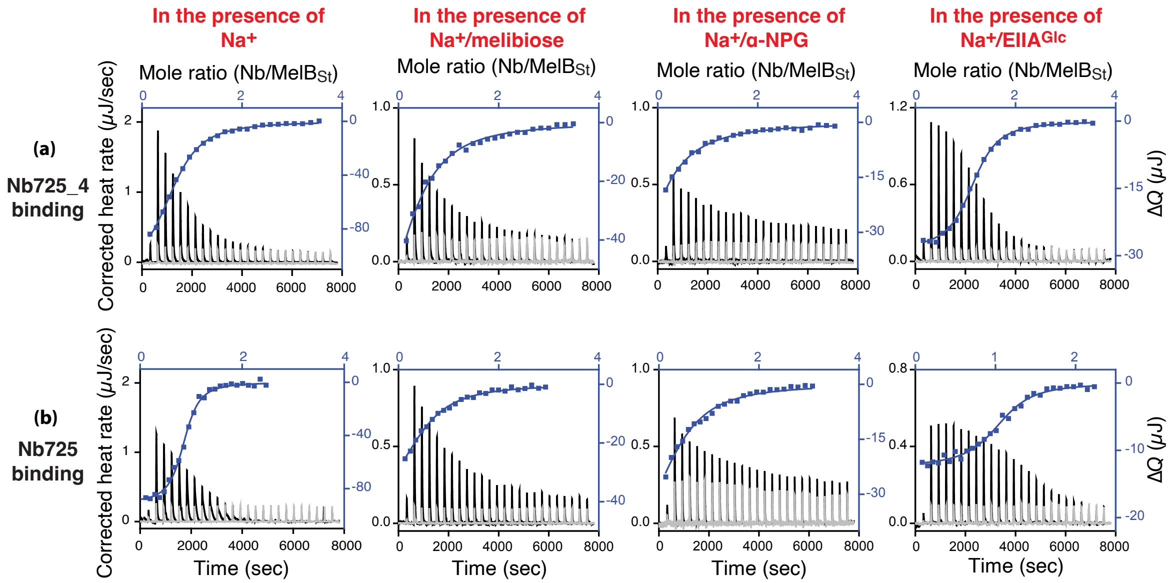

**sFig. 2. Effects of substrate/ligand binding on the Nbs binding.** All ITC binding measurements at 25 °C and curve fitting were described in the Methods. The purified Nb725_4, Nb725, $MelB_{St}$, and $EIIA^{Glc}$ protein samples were dialyzed against an identical buffer containing 100 mM NaCl. $MelB_{St}$ at a concentration of 35 μM was placed in the sample cells and titrated with Nb725_4 or Nb725, respectively. The thermograms of each titration (black) and the corresponding control by titrating Nbs into the protein-free buffer (light gray) were plotted by the bottom/left (x/y) axes. The binding isotherm and fitting using a one-site independent-binding model presented by top/right (x/y) axes. **(a)** Binding of Nb725_4 to the $Na^+$-bound $MelB_{St}$ in the absence or presence of melibiose (80 mM), α-NPG (5 mM), or $EIIA^{Glc}$ (2:1 ratio to $MelB_{St}$). **(b)** Binding of Nb725 to the $Na^+$-bound $MelB_{St}$ in the absence or presence of melibiose (80 mM), α-NPG (5 mM), or $EIIA^{Glc}$ (2:1 molar ratio to $MelB_{St}$).

**sFig. 3**

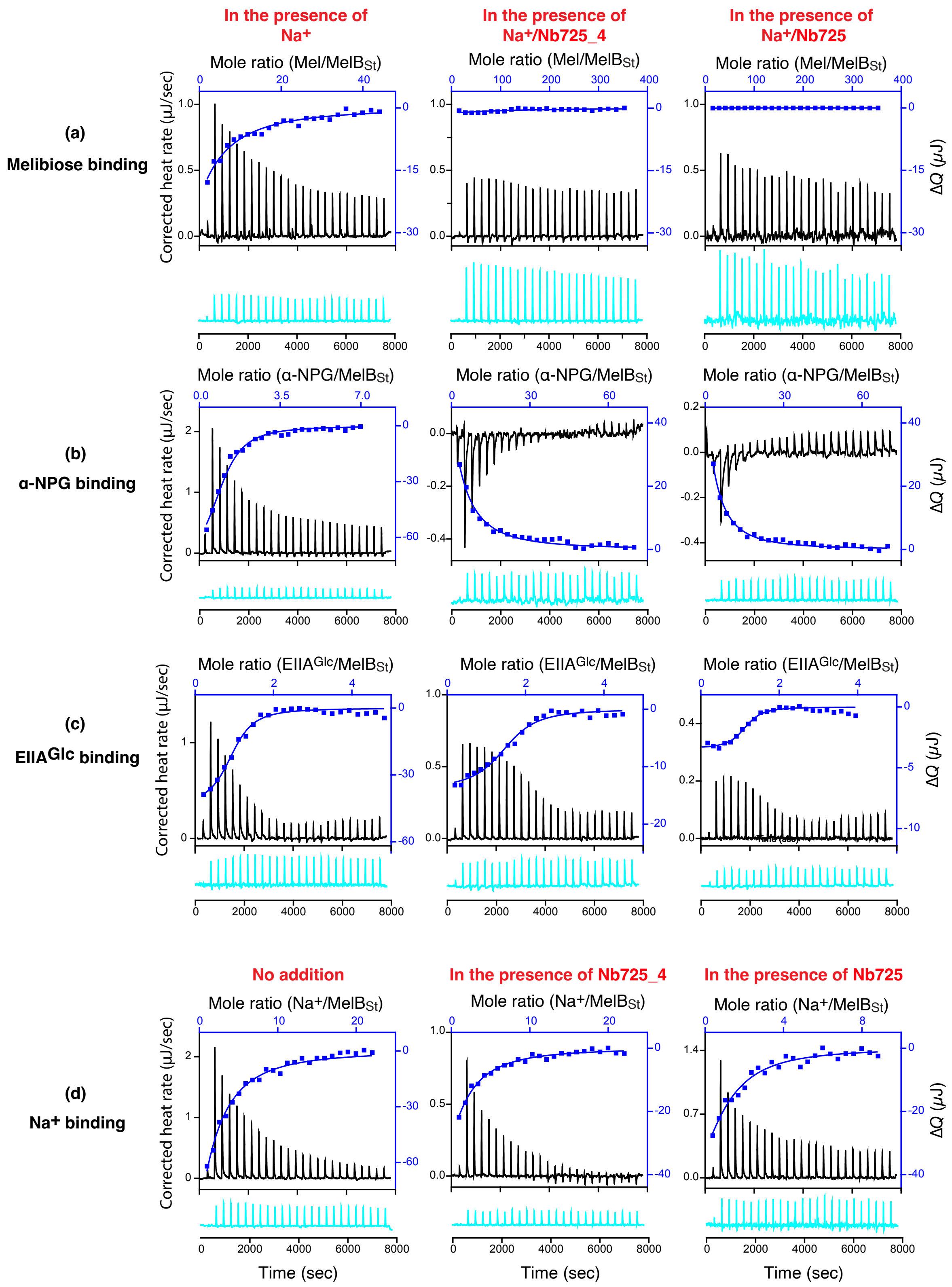

In the presence of Na+
In the presence of Na+/Nb725_4
In the presence of Na+/Nb725
(a) Melibiose binding
Mole ratio (Mel/MelBSt)
(b) α-NPG binding
Mole ratio (α-NPG/MelBSt)
(c) EIIAGlc binding
Mole ratio (EIIAGlc/MelBSt)
No addition
In the presence of Nb725_4
In the presence of Nb725
(d) Na+ binding
Mole ratio (Na+/MelBSt)
Corrected heat rate (μJ/sec)
ΔQ (μJ)
Time (sec)

**sFig. 3. Nb effects on substrate/ligand binding.** All ITC binding measurements at 25 °C and curve fitting were described in the Methods. The binding measurements of melibiose, α-NPG and $EIIA^{Glc}$ were conducted in buffer, 20 mM Tris-HCl, pH 7.5, 100 mM NaCl, 0.01% DDM, and 10% glycerol. The $Na^+$ binding measurements were conducted in 20 mM Tris-HCl, pH 7.5, 100 mM choline-Cl, 0.01% DDM, and 10% glycerol as described in (Hariharan, 2017). All proteins and ligand solutions were buffer-matched. $MelB_{St}$ complexed with Nb725_4 or Nb725 were prepared by mixing them at a 1:2 molar ratio. The thermograms of each titration (black) were plotted by bottom/left (x/y) axes and the binding isotherm and fitting using one-site independent-binding model presented by top/right (x/y) axes. The corresponding controls by titrating each ligand into the protein-free buffer (cyan) were presented below at an identical scale. **(a)** Titration of melibiose to $MelB_{St}$ or a given $MelB_{St}$/Nb complex at 80 μM for $MelB_{St}$. **(b)** Titration of α-NPG binding to $MelB_{St}$ a given $MelB_{St}$/Nb complex at 50 μM for $MelB_{St}$. **(c)** Titration of $Na^+$ to $MelB_{St}$ or a given $MelB_{St}$/Nb complex at 80 μM for $MelB_{St}$**. (d)** Titration of $EIIA^{Glc}$ to $MelB_{St}$ or a given $MelB_{St}$/Nb complex at 50 μM for $MelB_{St}$. For the titration into the $MelB_{St}$/Nb complexes, melibiose and α-NPG in the assay solutions were increased from 10 mM to 80 mM or 1 mM to 10 mM. The $Na^+$ at 3 mM and $EIIA^{Glc}$ at 0.5 mM were applied for titrations.

#### sFig. 4a

#### Overall strategy for single-particle resonstruction

**14,094 micrographs collected**
**13,649 curated**

**(a) Initial 3D reconstruction**

Template pick from 13,649 micrographs

2D Classification
Heterogeneous Refinement
3D Ab-inito Reconstruction
Non-uniformed Refinemnt

**41,264-particle set**

GSFSC resolution = 3.49 Å

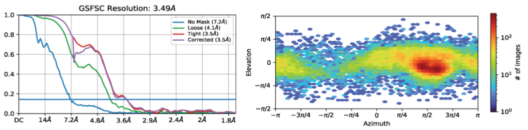

**(b) Map improvement**

Template pick from 13,649 micrographs

2D Classification
Heterogeneous Refinement
Local Refinement

Combinded/removed duplicates
Heterogeneous Refinement

**188,566-particle set**

GSFSC resolution = 3.58 Å

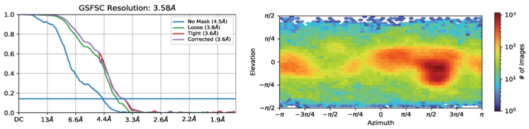

Re-cleaned particles focusing top/bottom views
from the rejected 2D classes

2D Classfication
Heterogeneous Refinement
Local Refinement

Combinded/removed duplicates
Heterogeneous Refinement
Local Refinement

**196,254-particle set**

GSFSC resolution = 3.39 Å

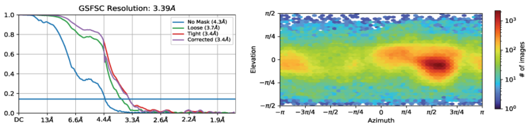

**(c) Tilted data collection**

**8,716 micrographs tilted collection at 30 °**
**7,129 curated**

Template pick from 7,129 micrographs

2D Classfication
Heterogeneous Refinement
Local Refinement

**182,101 particles selected**

Combinded
Heterogeneous Refinement
Local Refinement

**378,355-particle set**

GSFSC resolution = 3.40 Å

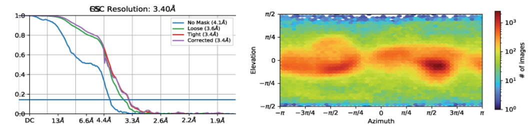

3D Classfication
Heterogeneous Refinement
Local Refinement

**296,925-particle set**

GSFSC resolution = 3.37 Å

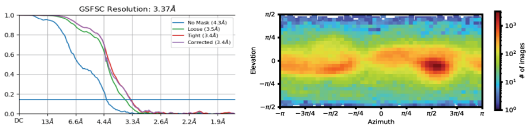

**sFig. 4b**

**Initial 3D reconstruction**

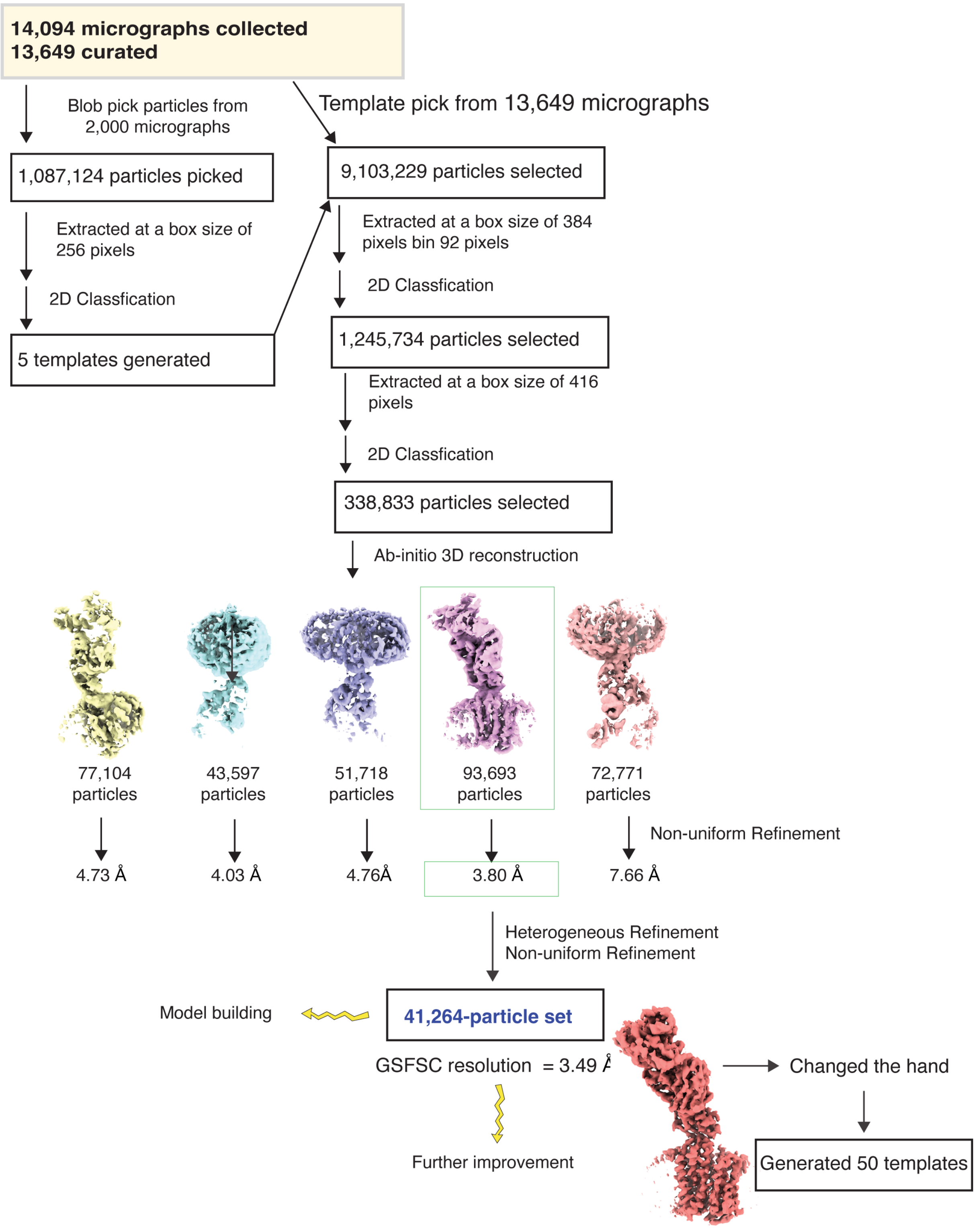

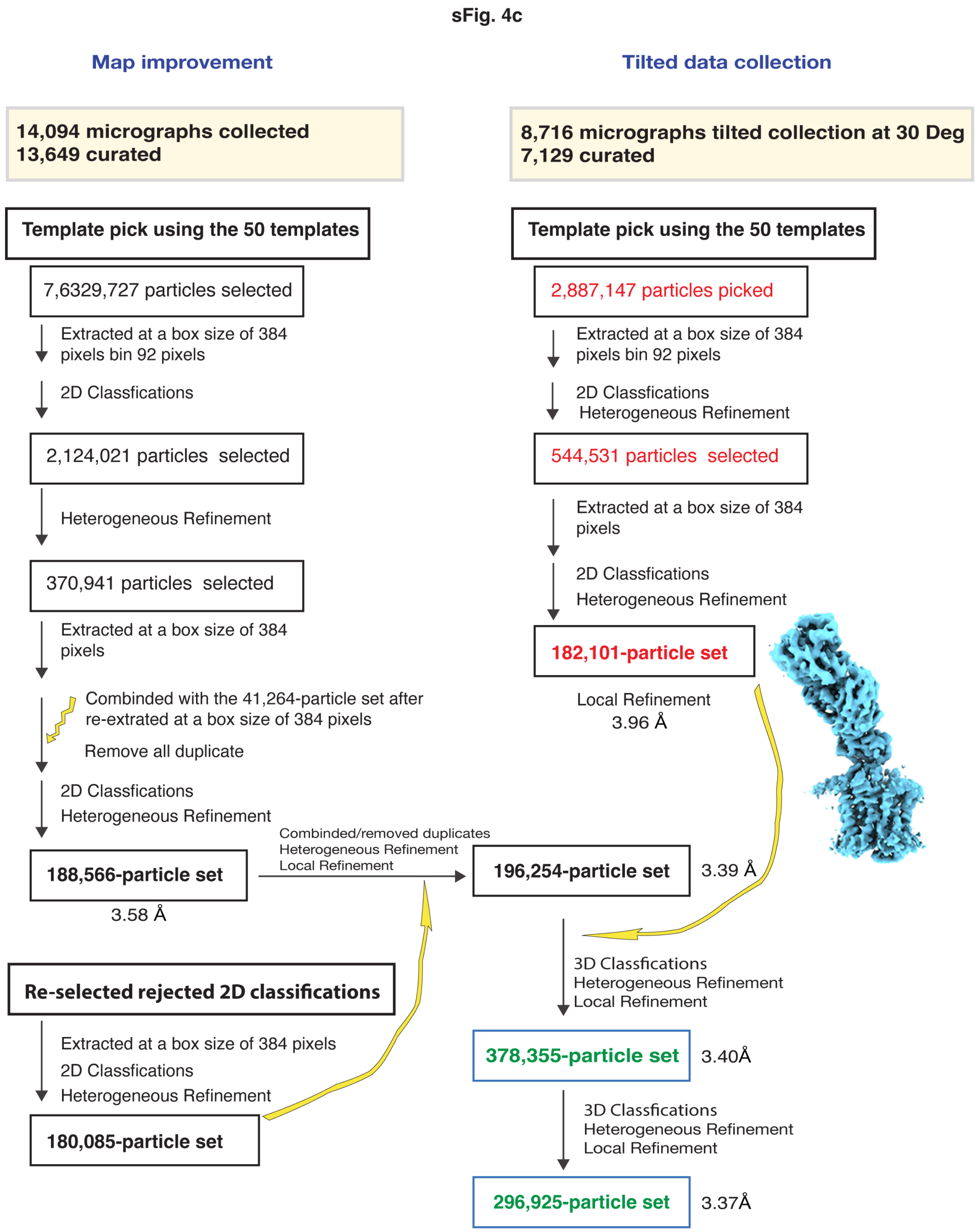

**sFig. 4. CryoEM data process.** The strategy for the reconstruction and refinement was outlined and the details were presented in panels b-c.

### sFig. 5

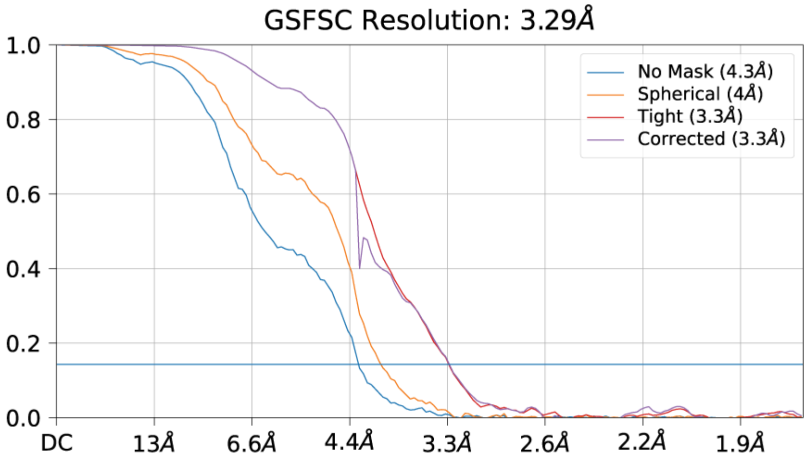

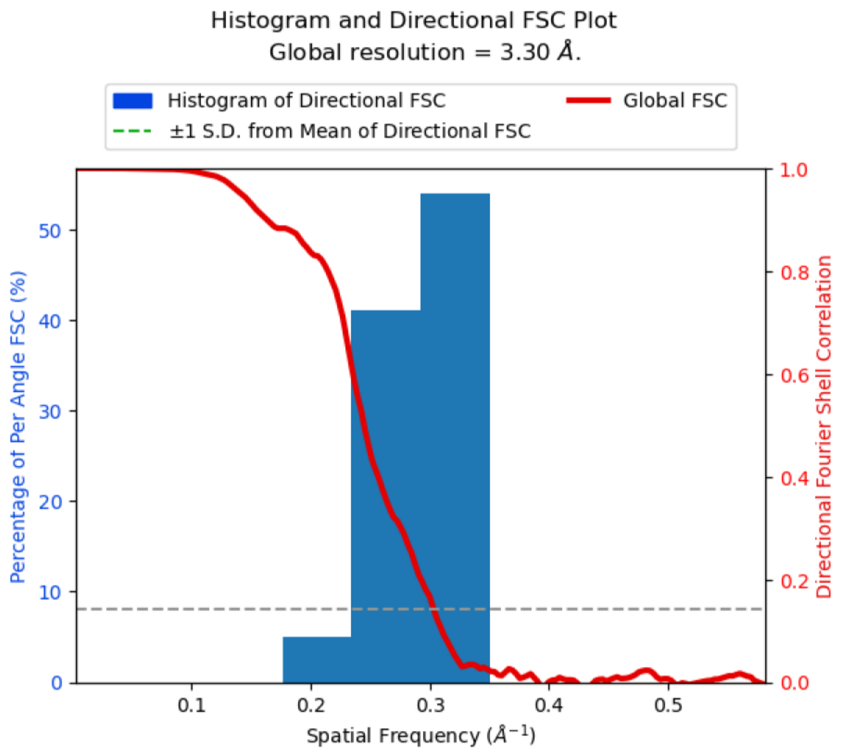

**sFig. 5. GSFSC resolution and 3dFSC.** The resolution of map and particle orientation distribution was assessed by cryoSPARC program using default setting parameters.

#### sFig. 6

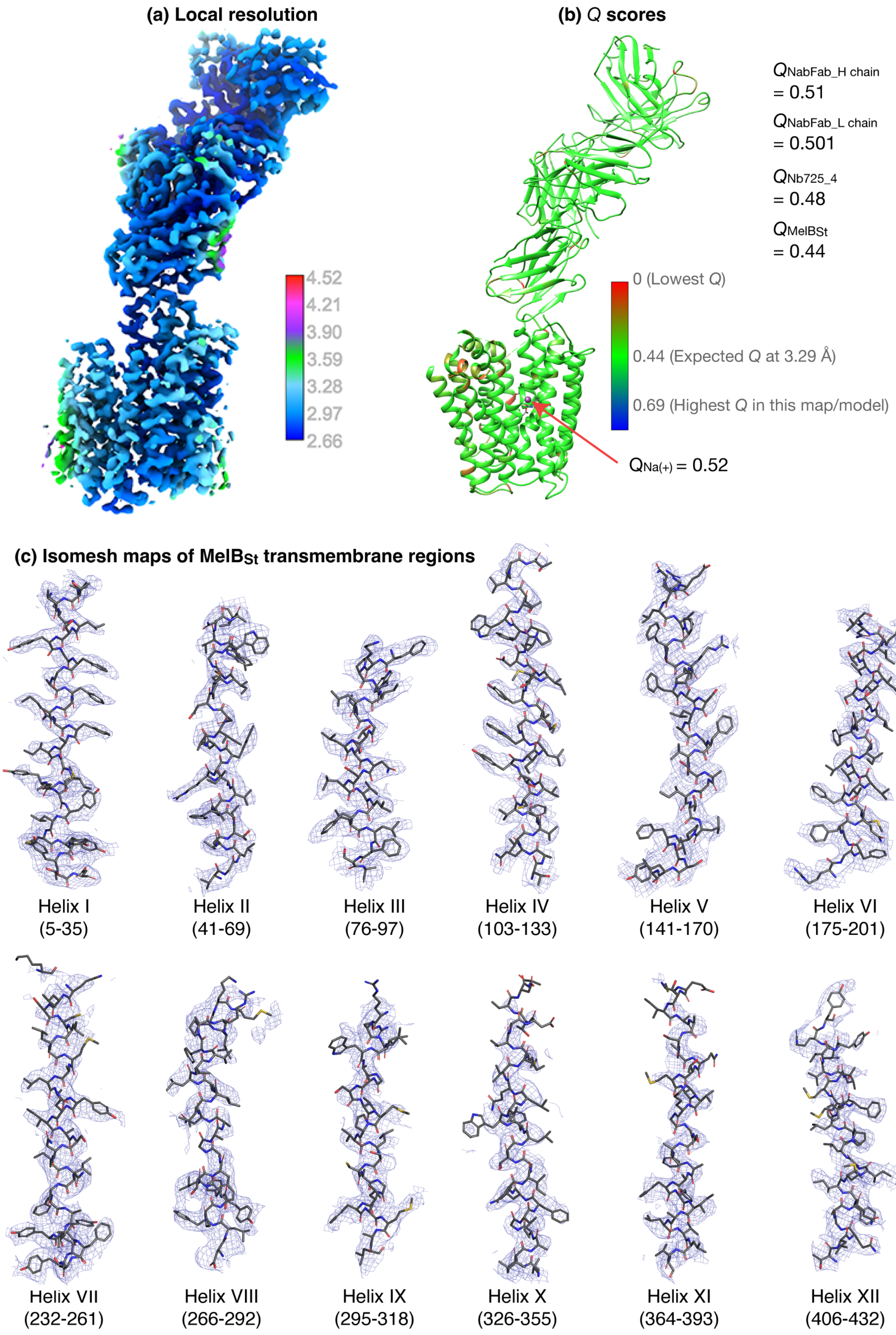
(a) Local resolution
4.52
4.21
3.90
3.59
3.28
2.97
2.66
(b) Q scores
QNabFab_H chain = 0.51
QNabFab_L chain = 0.501
QNb725_4 = 0.48
QMelBSt = 0.44
0 (Lowest Q)
0.44 (Expected Q at 3.29 Å)
0.69 (Highest Q in this map/model)
QNa(+) = 0.52
(c) Isomesh maps of MelBSt transmembrane regions
Helix I (5-35)
Helix II (41-69)
Helix III (76-97)
Helix IV (103-133)
Helix V (141-170)
Helix VI (175-201)
Helix VII (232-261)
Helix VIII (266-292)
Helix IX (295-318)
Helix X (326-355)
Helix XI (364-393)
Helix XII (406-432)

**sFig. 6. Evaluation of map and models. (a)** The local resolution. The map half_A and half_B files produced by the cryoSPARC Local Refinement program were used to calculate the Local Resolution Map by Phenix and displayed by UCSF ChimeraX using the defined color key. **(b)** *Q* scores. The auto-sharpened map generated by Phenix under the default setting was used as the main map for the model building and also for *Q*-score calculations by Map*Q* program in UCSF Chimera against the final structure at a sigma level of 0.4. The results of the *Q* scores for each residue were color-painted on the structure using the given color key. The bound $Na^+$ was shown as a sphere. The expected score at a resolution of 3.29 Å is 0.44. **(c)** Density mapped on the $MelB_{St}$ transmembrane helices. The auto-sharped map by Phenix was used to calculate the isomesh maps for each transmembrane helix at a sigma level of 10 and carving of 1.8.

sFig. 7

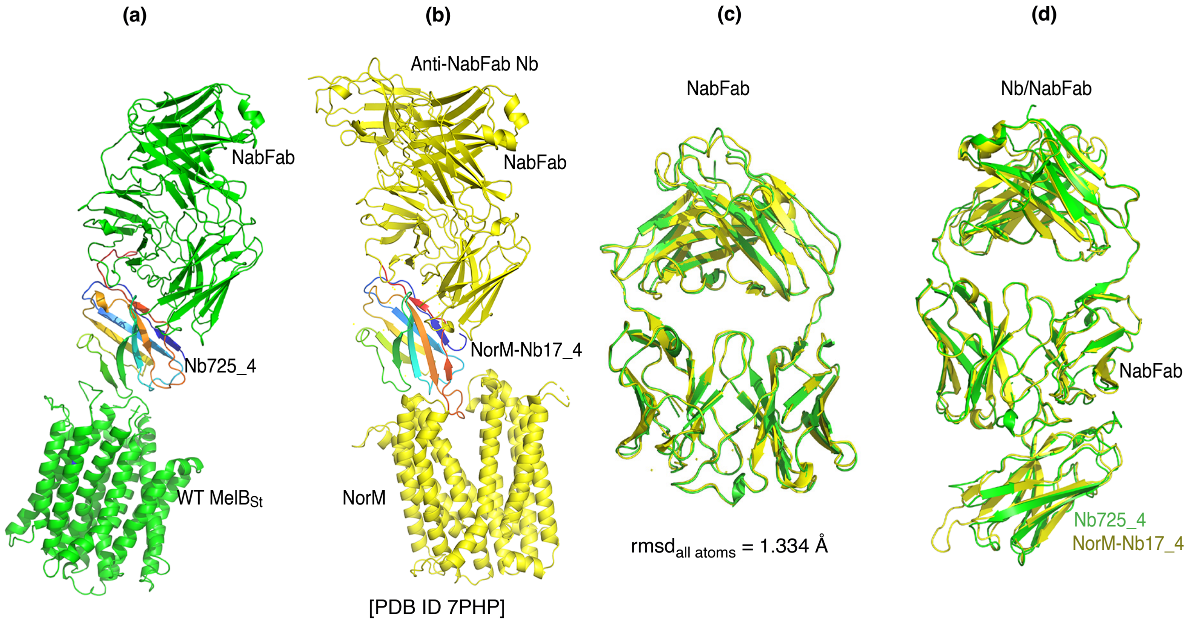

**sFig. 7. NabFab comparison.** (**a & b**) NorM complexed with anti-NorM Nb17_4/NabFab/anti-NabFab Nb [PDB ID 7PHP] was superimposed with $MelB_{St}$ complexed with Nb725-4 and NabFab based on the H chains of NabFab in the two structures. Nb725-4 and NorM-Nb17_4 were colored in the rainbow. (**c**) The NabFab in the NorM complex [PDB ID 7PHP] was superimposed with that in $MelB_{St}$ complex. (**d**) The NabFab/Nb in the NorM complex was superimposed with that in $MelB_{St}$ complex based on their H chains. Green, $MelB_{St}$ complex; yellow, NorM complex.

**sFig. 8**

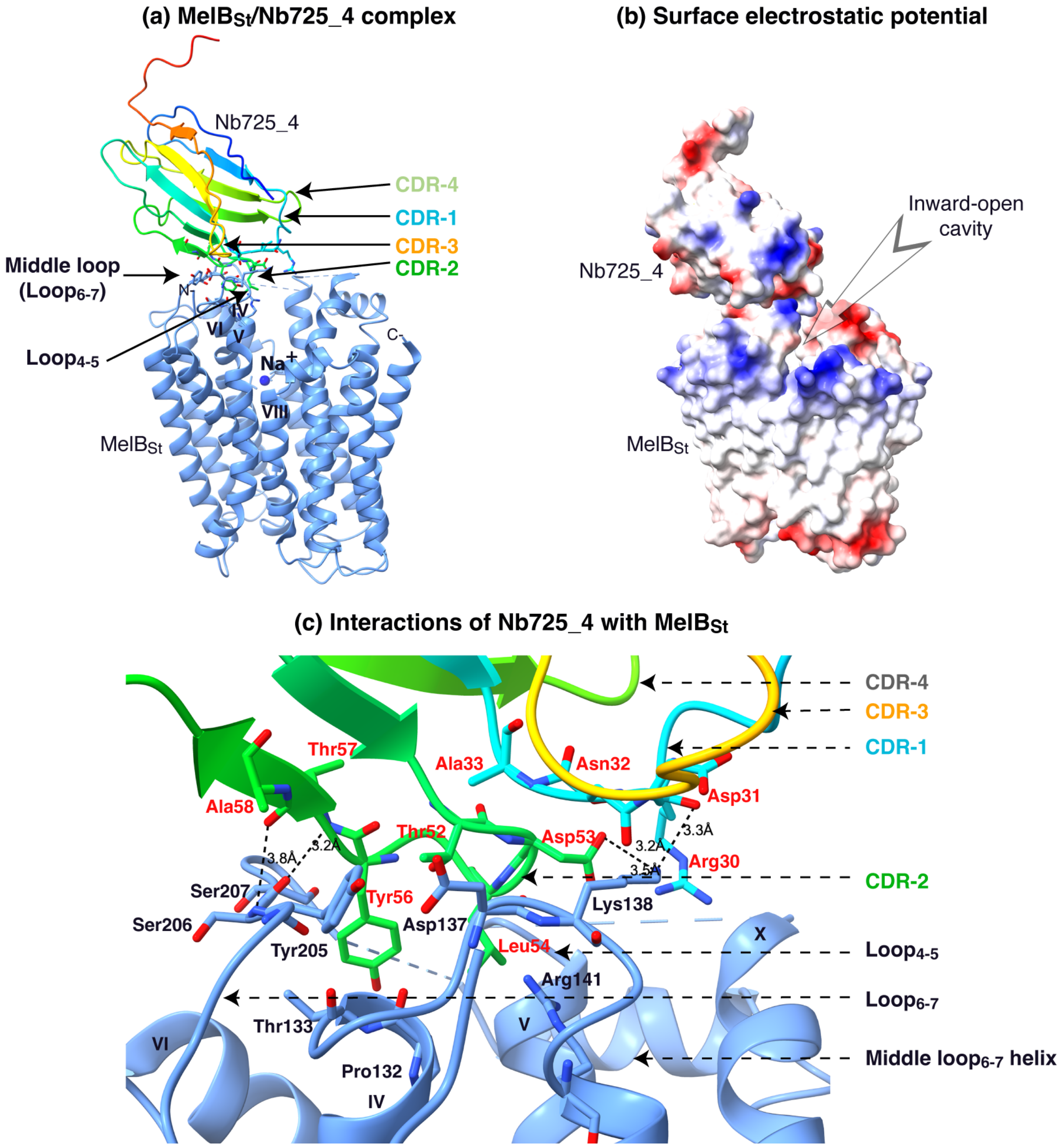

**sFig. 8. Interactions of Nb725m_4 and MelB$_{St}$. (a)** Nb725_4 bound to the N-terminal domain of MelB$_{St}$. The contact between Nb CDR-1 and CDR-2 and MelB$_{St}$ loop$_{4-5}$, and loop$_{6-7}$ contributed to the major interactions. **(b)** The surface electro potential map was generated in the UCSF Chimera X program. The inward-open cavity is indicated. **(c)** The binding interface. All residues with a buried solvent-accessible surface area >15 Å$^2$ or polar interaction at a distance <3.5 Å were selected by the UCSF Chimera X program were highlighted in sticks. The salt-bridging and hydrogen-bonding interactions were highlighted by the dashed line. The Nb CDRs and epitope are indicated.

sFig. 9

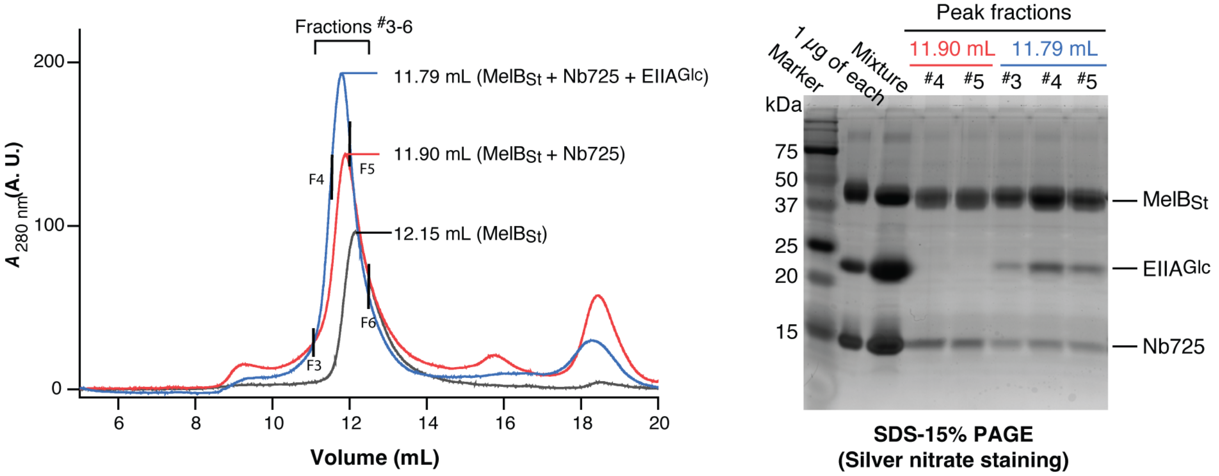

**sFig. 9. Complex of MelB$_{St}$ with Nb725 and EIIA$^{Glc}$.** Samples containing MelB$_{St}$ (black curve), MelB$_{St}$ with Nb725 (red curve), or with Nb725 and EIIA$^{Glc}$ (blue curve), were prepared and analyzed by gel filtration chromatography in a combination of the SDS-15%PAGE stained by silver nitrate. Mixture, the solution was prepared prior to the gel filtration chromatography containing all three proteins. 1 μg of each protein was loaded as the control.

sFig. 10

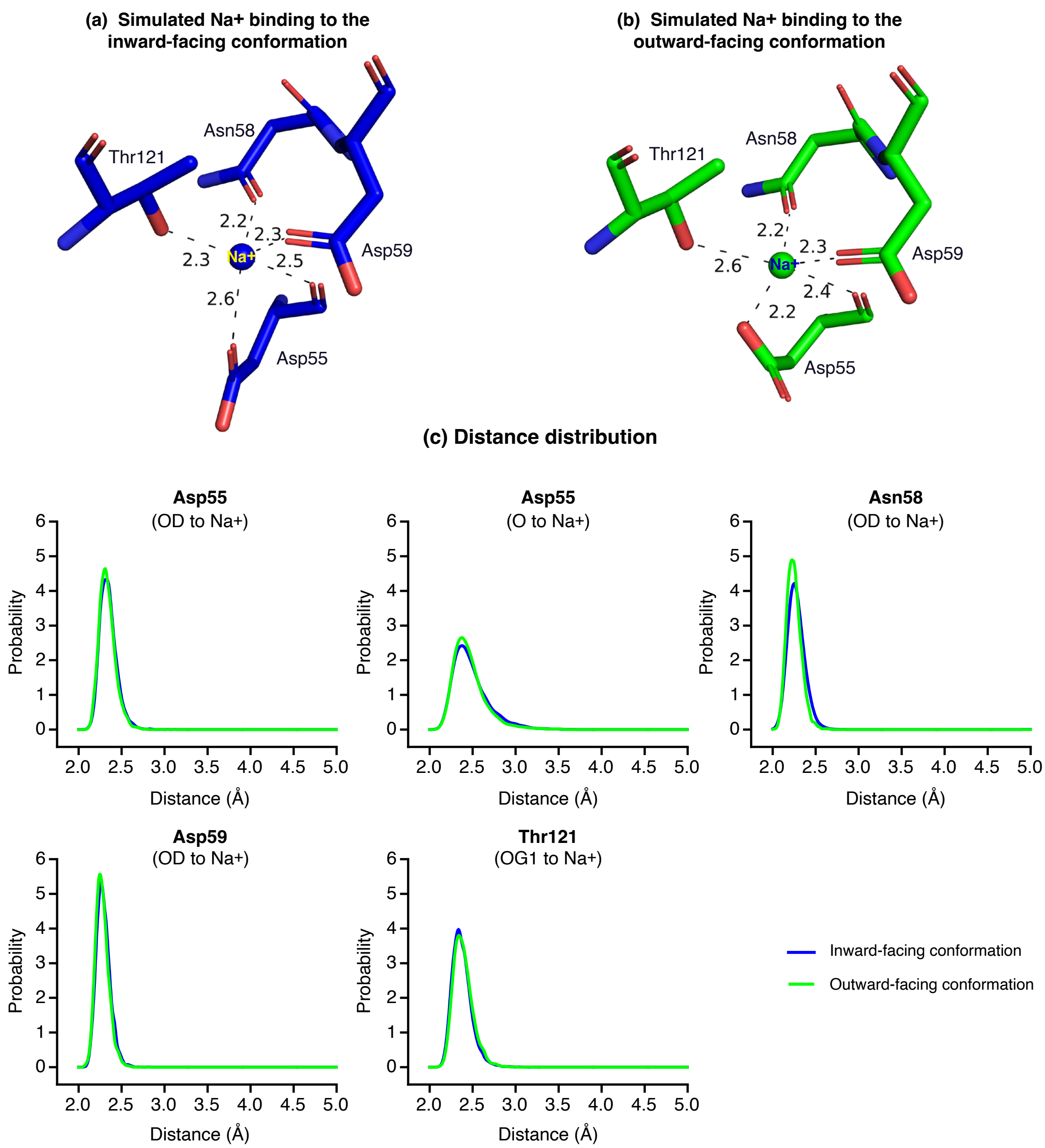

**sFig. 10. MD simulations of the Na$^+$ binding at both inward- and outward-facing states.** The equilibrated structures of Na$^+$-binding site for the inward- and outward-facing MelB$_{St}$ in the absence of melibiose. Distances between Na$^+$ and nearby residues' coordinating atoms are labeled in text and dashed lines. **(a)** Na$^+$ binding in the inward-facing conformation. **(b)** Na$^+$ binding in the outward-facing conformation. **(c)** Probability distribution of the distance between the bound Na$^+$ to all ligands. Blue, inward-facing conformation; green, outward-facing conformation.

**sFig. 11**

**Outward-facing X-ray crystal structure**

[PDB ID 7L17; 3.05 Å]

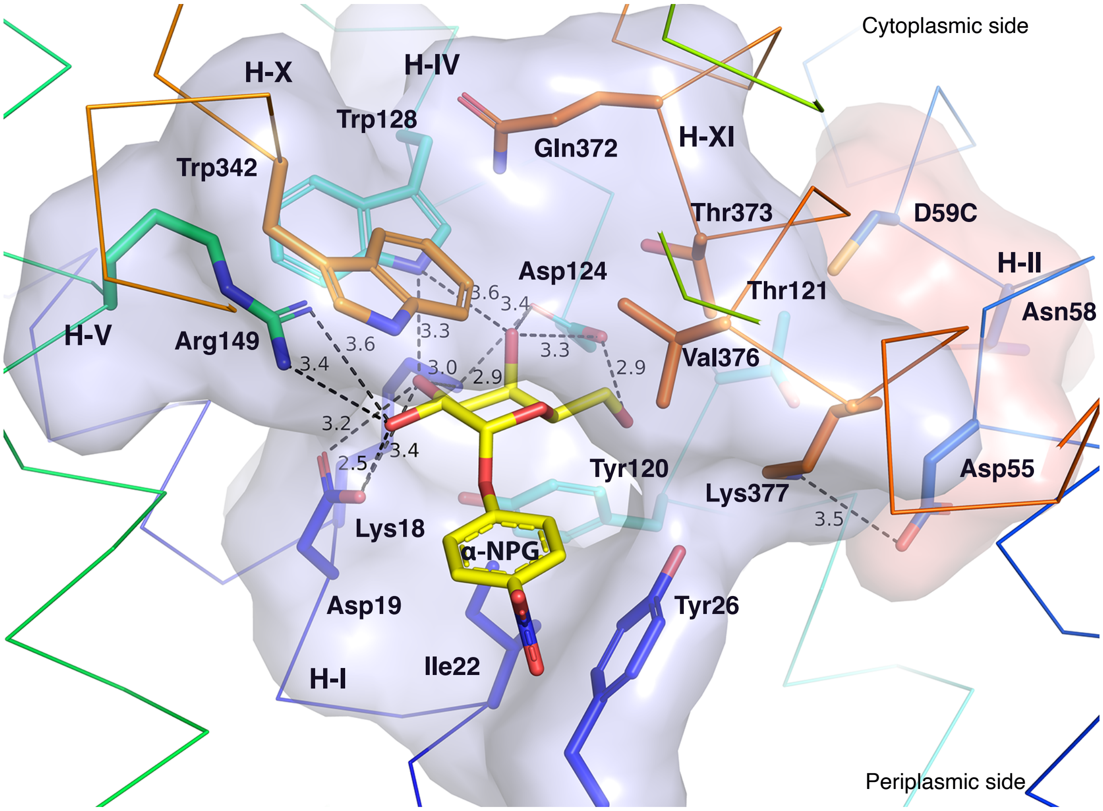

**sFig. 11. Galactose-binding pocket in the outward-facing crystal structure [PDB ID 7L17].** All polar interactions between the bound α-NPG and D59C $MelB_{St}$ were indicated by dash lines and the pocket was shown in surface representation in blue. The $Na^{+}$-binding residues were highlighted by sticks and surface representation in pink. The cytoplasmic and periplasmic sides were indicated. Eight N-terminal residues (Lys18, Asp19, Ile22, Tyr26, Tyr120, Asp124, Trp128, and Arg149), especially the four charged residues, form the multiple polar interactions with one surface of the galactopyranosyl moiety. Five C-terminal residues (Trp342, Gln372, Thr373, Val376, and Lys377) form a non-specific barrier without a strong polar interaction

**sFig. 12**

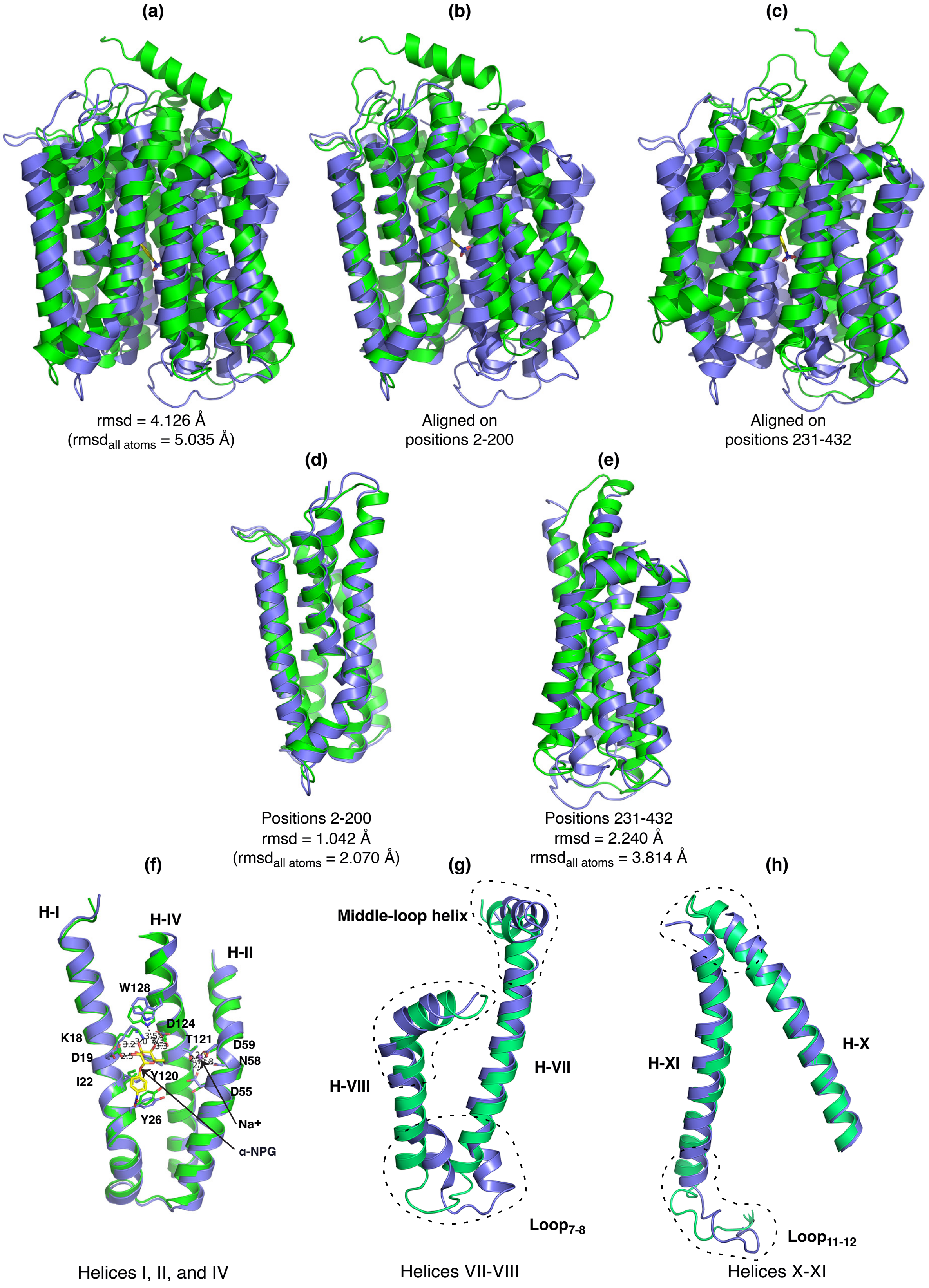

(a)
(b)
(c)
rmsd = 4.126 Å
(rmsd$_{\text{all atoms}}$ = 5.035 Å)
Aligned on
positions 2-200
Aligned on
positions 231-432
(d)
(e)
Positions 2-200
rmsd = 1.042 Å
(rmsd$_{\text{all atoms}}$ = 2.070 Å)
Positions 231-432
rmsd = 2.240 Å
rmsd$_{\text{all atoms}}$ = 3.814 Å
(f)
(g)
(h)
H-I
H-IV
H-II
W128
D124
K18
T121
D59
D19
N58
I22
Y120
D55
Y26
Na+
α-NPG
Middle-loop helix
H-VIII
H-VII
Loop$_{7\text{-}8}$
H-XI
H-X
Loop$_{11\text{-}12}$
Helices I, II, and IV
Helices VII-VIII
Helices X-XI

**sFig. 12. Alignment.** The alignment of the sugar-bound outward-facing structure (7L17, green) with the $Na^+$-bound inward-facing structure (blue) were carried out in Pymol or Coot programs. The rmsd values reported from Pymol program were either based on default settings with outlier rejection or using all atoms after removing unmatched residues. **(a)** Superposition of both structures. (**b**) Full-length alignment based on positions 2-200. **(c)** Full-length alignment based on positions 231-332. (**d)** Focused alignment of positions 2-200. **(e)** Focused alignment of positions 231-432. **(f)** Helices I, II, and IV isolated from the focused alignment of positions 2-300 in *d*. The major binding residues for $Na^+$ and α-NPG were highlighted. **(g)** Helices VII-VIII and the extended loops isolated from the focused alignment of positions 219-432. **(h)** Helices X-XI and the extended loops isolated from the focused alignment of positions 219-432.

sFig. 13

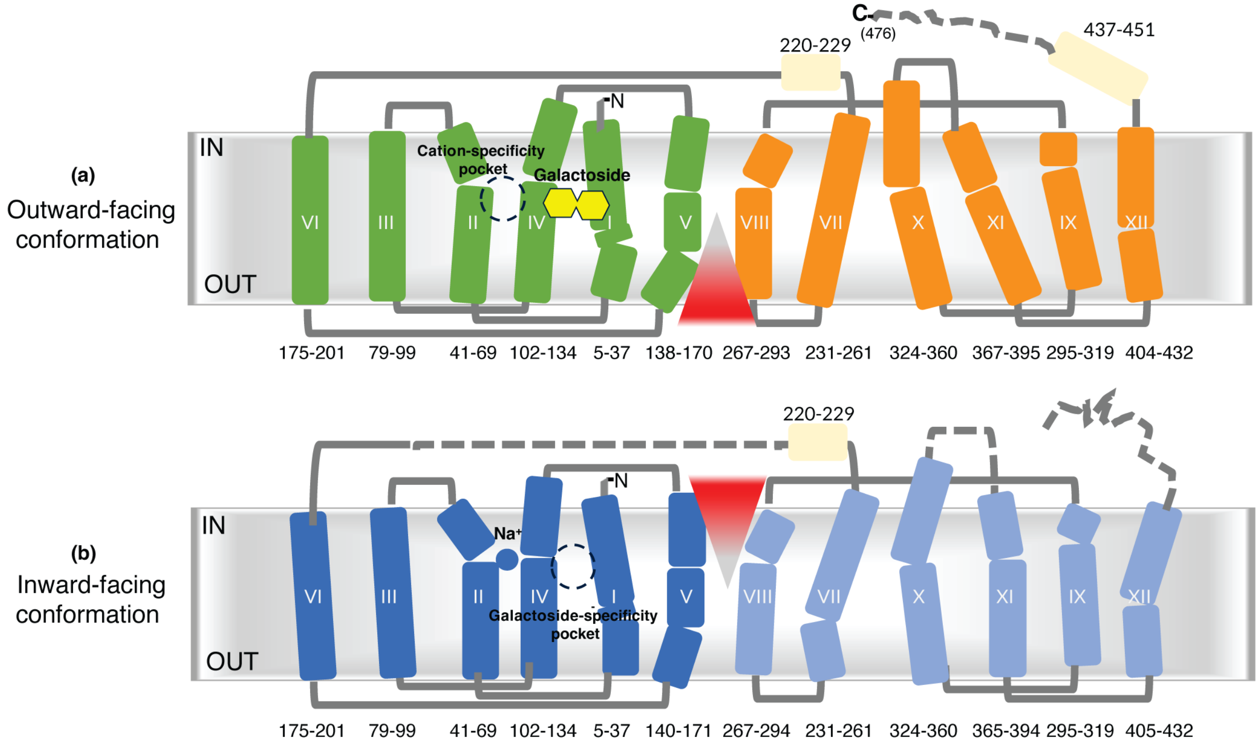

**sFig. 13. Membrane topology. (a)** The sugar-bound outward-facing structure (PDB ID 7L17). **(b)** The $Na^+$-bound inward-facing structure. The full-length $MelB_{St}$ is illustrated by transmembrane topology based on both structures. The residue positions for each helix are indicated. The N- and C-terminal transmembrane helices are colored green and orange for the outward-facing conformation, respectively, as well as blue and light blue for the inward-facing conformation, respectively. The peripheral helices are colored in light yellow. Blue ball, $Na^+$; yellow hexagons, galactoside. The red triangle indicates the solvent-access paths. Grey lines, loops; dashed lines, un-resoled loops. Broken circles, either cation-specificity pocket or sugar-specificity pocket.

sFig. 14

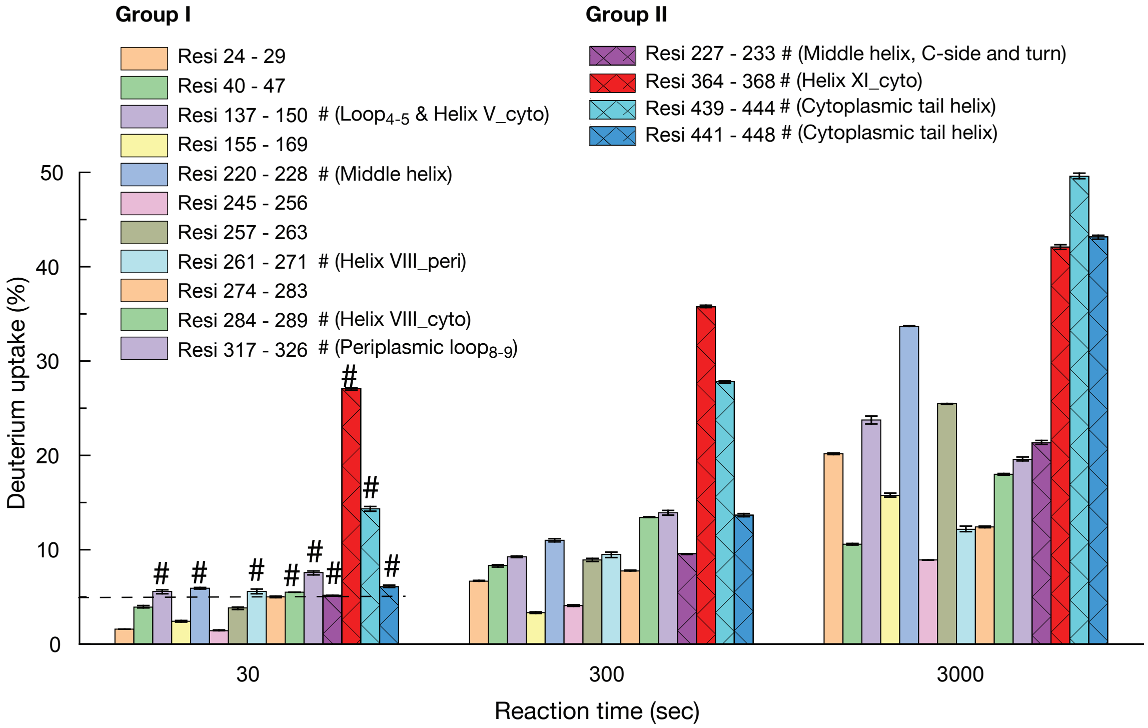

**sFig. 14. Histograms of deuterium uptake time courses.** The deuterium uptake time courses of the groups I&II presented in the Fig. 7 were replotted in the histogram. Error bar, sem; test number, 3.
