## Extended figure for "Mobile barrier mechanisms for Na^+^-coupled symport in an MFS sugar transporter"

**Extended Fig. 1**

**Deuterium uptake time-course of all peptides with statastically significant changes**

**(Coverage of residue positions 2 - 263)**

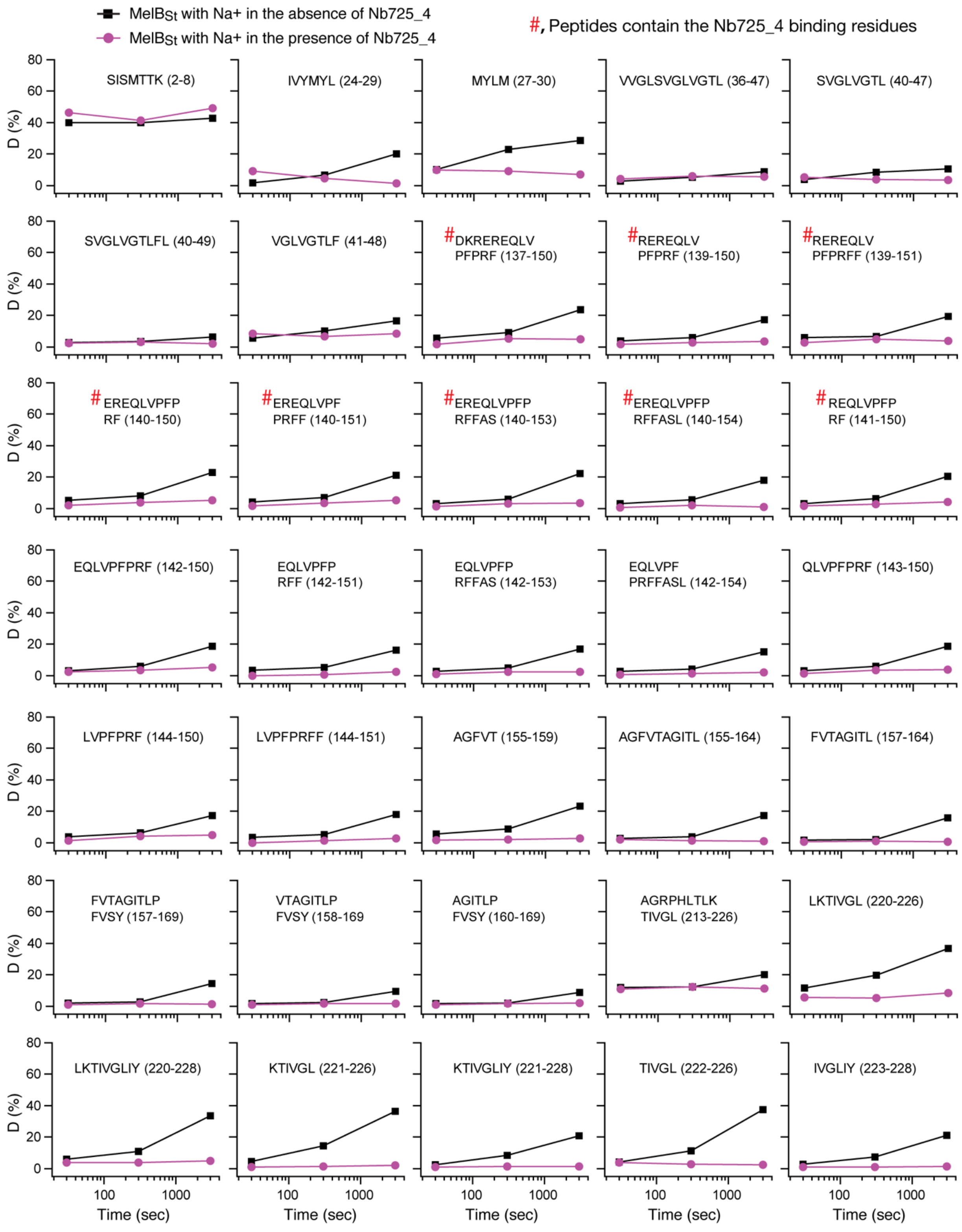

(Coverage of residue positions 261 - 475)

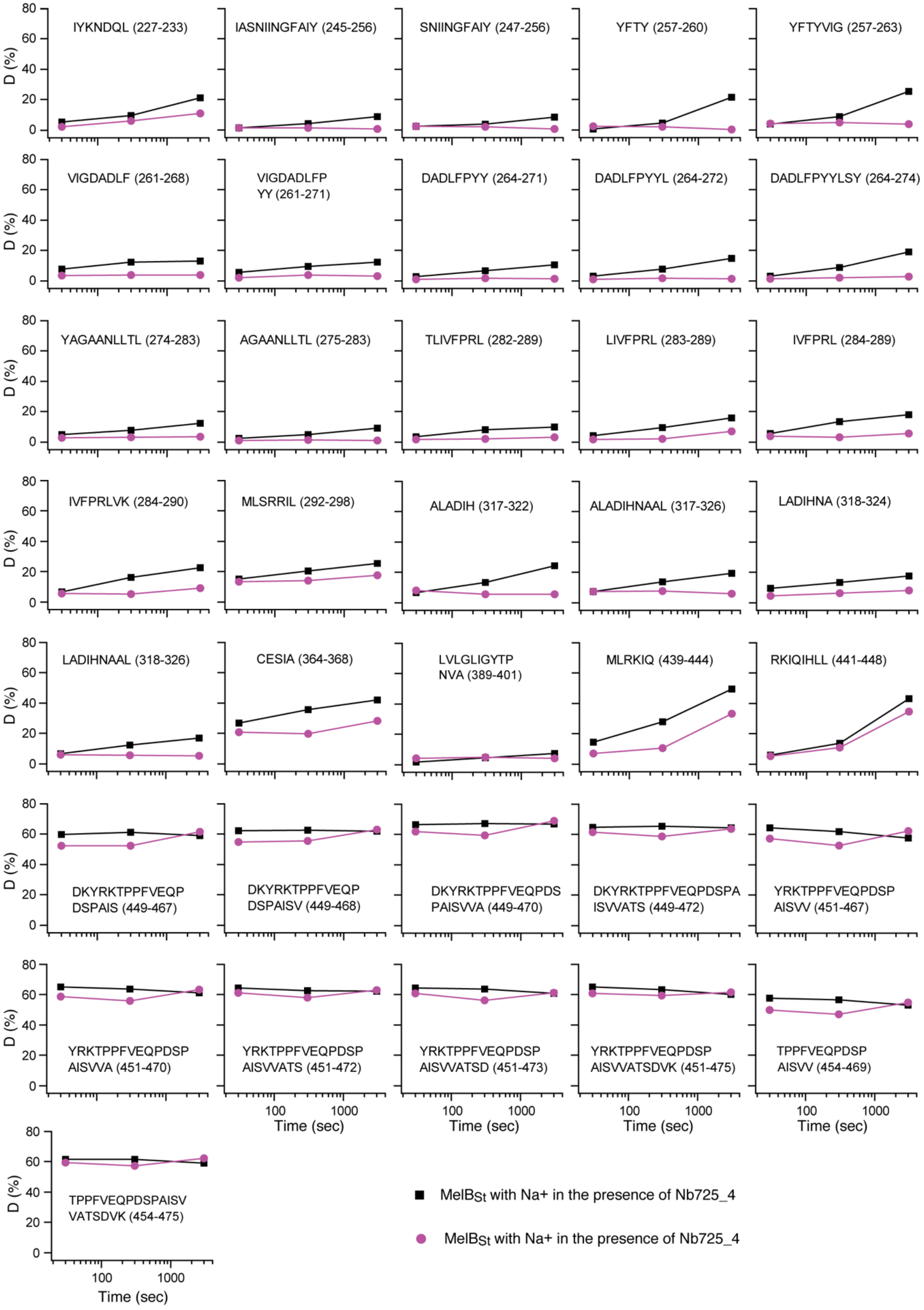
